# Transient response of basal ganglia network in healthy and low-dopamine state

**DOI:** 10.1101/2021.01.05.425413

**Authors:** Kingshuk Chakravarty, Sangheeta Roy, Aniruddha Sinha, Atsushi Nambu, Satomi Chiken, Jeanette Hellgren Kotaleski, Arvind Kumar

## Abstract

The basal ganglia (BG) are crucial for a variety of motor and cognitive functions. Changes induced by persistent low-dopamine (e.g. in Parkinson’s disease), result in aberrant changes in steady-state population activity (*β*-band oscillations) and transient response of the BG. Typically, brief cortical stimulation results in a triphasic response in the substantia nigra pars reticulata (SNr, output of the BG). The properties of the triphasic responses are shaped by dopamine levels. While it is relatively well understood how changes in BG result in aberrant steady state activity, it is not clear which BG interactions are crucial for the aberrant transient responses in the BG. Moreover, it is also not clear whether the same or different mechanisms underlie the aberrant changes in steady-state activity and aberrant transient response. Here we used numerical simulations of a network model of BG to identify the key factors that determine the shape of the transient responses. We show that an aberrant transient response of the SNr in low-dopamine state, involves changes in both, the direct pathway and the recurrent interactions within the globus pallidus externa (GPe) and between GPe and sub-thalamic nucleus. We found that the connections from D2-type spiny projection neurons to GPe are most crucial in shaping the transient response and by restoring them to their healthy level, we could restore the shape of transient response even in low-dopamine state. Finally, we show that the changes in BG that result in aberrant transient response are also sufficient to generate pathological oscillatory activity.

**Significance statement:** To understand how changes induced by low-dopamine (e.g. in Parkinson’s disease, PD) affect basal ganglia (BG) function, we need to identify the factors that determine the shape of BG responses to brief cortical stimuli. We show that transient response of the BG is also affected by recurrent interactions within the subnuclei of the BG, and not just feedforward pathways. We found that input and local connectivity within the globus pallidus externa are crucial for shaping the transient response. We also show that the same network changes may underlie both, pathological *β*-band oscillations and aberrant transient responses. Our results highlight the importance of the recurrent connectivity within the BG and provide a coherent view of emergence of pathological activity in PD.

## Introduction

Parkinson’s disease (PD) is a debilitating neurodegenerative brain disease with multiple cognitive and motor symptoms. Etiologically the disease is attributed to the progressive loss of dopaminergic neurons in the substantia nigra pars compacta. Dopamine affects neuronal excitability, synaptic strength and synaptic plasticity. Consistent with this, data from human patients and animal models show that dopamine deficit results in a number of changes in the neuronal activity especially in the basal ganglia (BG). At the level of neuronal activity, in PD, synchronized *β*-band oscillations (15-30 Hz) bursts in the globus pallidus externa (GPe) and sub-thalamic nucleus (STN) (Brown et al., 2001; Tinkhauser et al., 2017; Raz et al., 2000; Mallet et al., 2008) emerge along with an increase in spike bursts (Tachibana et al., 2011; Nambu et al., 2015). In the striatum, firing rate of D2-type dopamine receptor expressing spiny projection neuron (D2-SPN) is increased whereas firing rate of D1-SPNs is reduced (Mallet et al., 2006; Sharott et al., 2017). Moreover, while cortical inputs to D2-SPN are enhanced, inputs to D1-SPN are weakened (Ketzef et al., 2017; Parker et al., 2016; Filipović et al., 2019). The aforementioned changes in the activity and structure of the BG are persistent and indicate a change2016 in ‘operating point’ of the BG. But these observations do not provide mechanistic links between behavior deficits of PD and BG activity.

During action-selection or decision-making tasks the BG receive transient inputs (Gage et al., 2010) from different cortical regions. It is therefore, important to understand how the response of the BG network to a transient cortical input is altered during PD condition. In a healthy state, transient cortical stimulation elicits a triphasic response (composed of early excitation, inhibition, and late excitation) at the population level in the output nuclei of the BG i.e. globus pallidus interna (GPi) or substantia nigra pars reticulata (SNr) (Sano et al., 2013; Chiken and Nambu, 2013; Ozaki et al., 2017). The triphasic response is consistent with the predictions of a simple feedforward model of the BG involving the so-called direct, indirect and hyper-direct pathway (Albin et al., 1989; Jaeger and Kita, 2011). However, individual neurons in SNr (Sano and Nambu, 2019) or GPi (Iwamuro et al., 2017) can show biphasic or monophasic responses. In dopamine depleted conditions, the fraction of neurons showing biphasic and monophasic responses is changed resulting in an altered population response.

To identify what determines the shape of BG transient responses we used a computational model of the BG (Lindahl and Kotaleski, 2016). We found that, consistent with experimental data (Sano and Nambu, 2019) and predictions of the feedforward model of the BG (Albin et al., 1989), in healthy state, the SNr showed triphasic shaped responses for brief cortical inputs. In the low-dopamine state, with the default settings, the SNr transient response was biphasic. However, by changing the strength of synapses along the direct and indirect pathways (D2-SPN→GPe-TI, and GPe-TI→STN) it was possible to observe the triphasic responses even in low-dopamine state. Interestingly, we found that changes in the transient response properties in PD state involve not only changes in the feed-forward connections (e.g. D1-SPN→SNr) but also recurrent interactions within BG subnuclei, e.g. the recurrent connections within the GPe (GPe-TA↔GPe-TI) and between GPe and STN (GPe*↔*STN). Next, we show that by restoring the connection from D2-SPN to GPe (D2-SPN→GPe-TI) to a normal value, even in low-dopamine state we can recover a transient response similar to that observed in healthy state. Thus, the D2-SPN→GPe-TI emerged as the most important descriptor of the aberrant transient response. Interestingly, it is the same connection that can unleash *β*-band oscillations (Kumar et al., 2011; Mirzaei et al., 2017). Thus, the same changes that underlie the emergence of pathological *β*-band oscillations, also make the transient response pathological. Thus, our results highlight the importance of the recurrent connections within the BG for the processing of transient information and lead to testable predictions.

## Material and Methods

### Neuron model

In order to achieve a good trade-off between simulation efficacy and ability to capture the neuronal dynamics, we used two types of neuron models in our BG network. Striatal D1 and D2 type dopamine receptor expressing spiny neurons (D1-SPN and D2-SPN), fast spiking interneurons (FSI) and STN neurons were realized using the standard leaky-integrate-fire neuron (LIF) model. The subthreshold dynamics of the membrane potential *V*^*x*^(*t*) was governed by the equation 1:

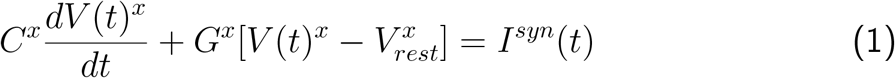

where *x* ∈ {D1-SPN, D2-SPN, STN}. In the equation 1, *C*^*x*^, *G*^*x*^, *V*_*rest*_ represent membrane capacitance, leak conductance and resting potentials, respectively. When *V*^*x*^ reaches the threshold potential 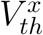, a spike is elicited and *V*^*x*^ is reset to 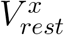 for refractory duration *t*_*ref*_ = 2 ms. *I*^*syn*^(*t*) models the total synaptic input current received by the neuron (see Figure 1 for the various sources of inputs to these neurons).

**Figure 1:**
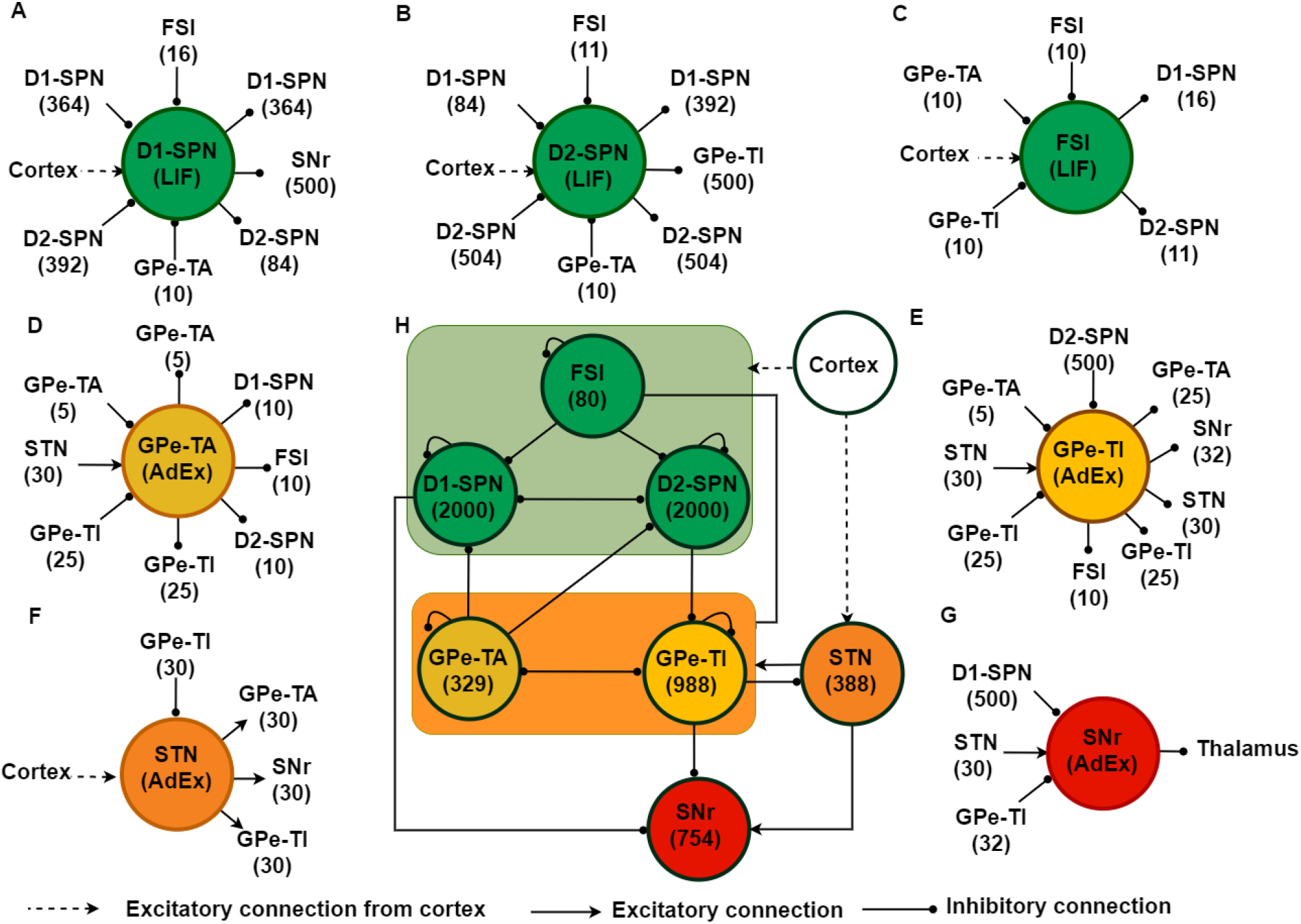
Schematic of the basal ganglia (BG) network model. (**A-G**) Schematic description of total number of inputs and outputs of a typical neuron in different subnetworks of the BG. (**H**), BG network structure along with the population size of individual nucleus. Within the BG network, the solid black lines with a circle at the end represent inhibitory synaptic connections and solid arrow lines represent excitatory synaptic connections. Dashed arrows denote the cortical excitatory input to BG.

All the parameter values for D1-SPN, D2-SPN, FSI and STN are summarized in the Table 3, 4, 5, and 8, respectively.

**Table 3:**
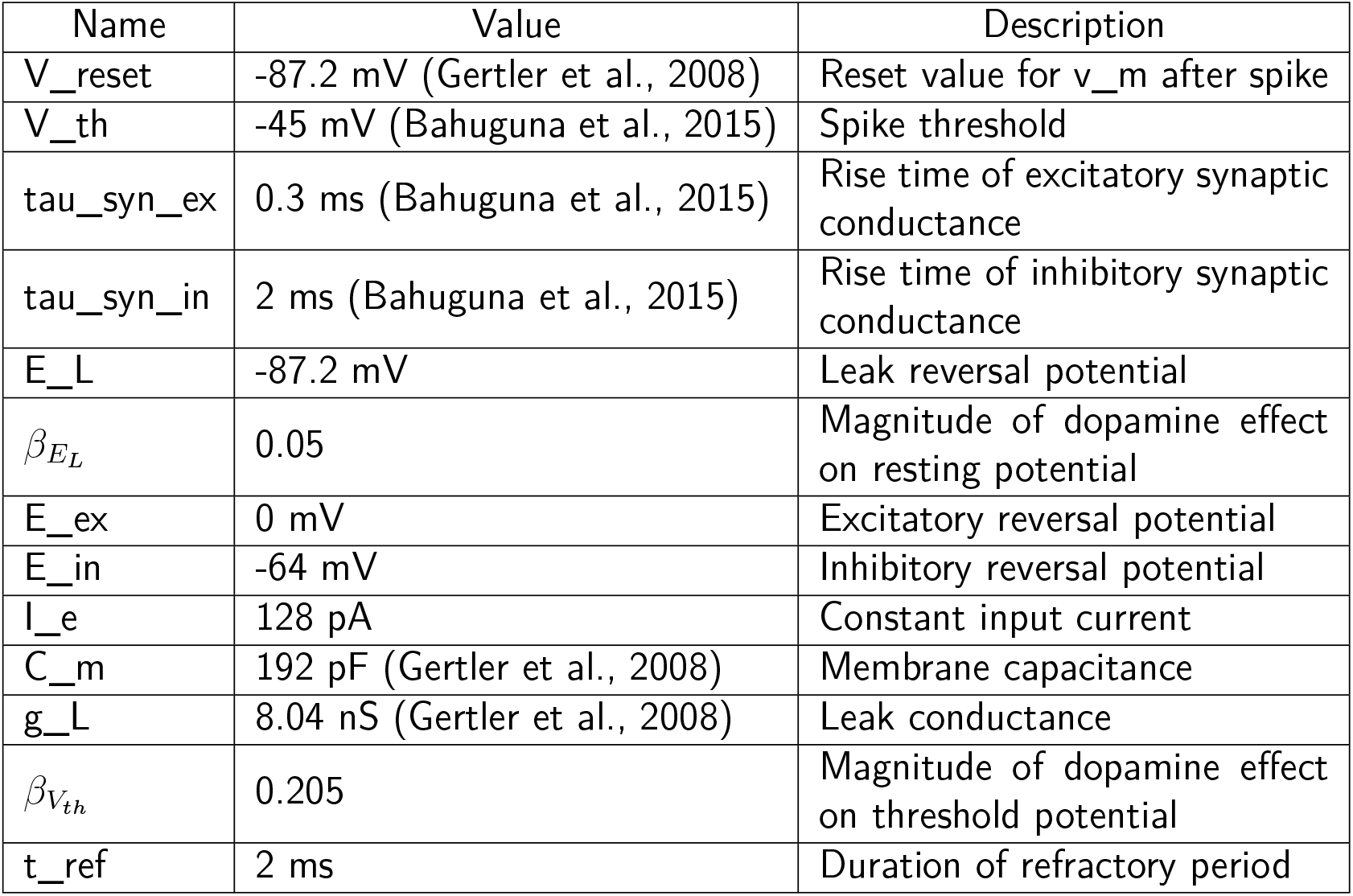
D1-SPN neuron parameters (leaky integrate and fire model).

| Name | Value | Description |
| --- | --- | --- |
| V_reset | -87.2 mV (Gertler et al., 2008) | Reset value for v_m after spike |
| V_th | -45 mV (Bahuguna et al., 2015) | Spike threshold |
| tau_syn_ex | 0.3 ms (Bahuguna et al., 2015) | Rise time of excitatory synaptic conductance |
| tau_syn_in | 2 ms (Bahuguna et al., 2015) | Rise time of inhibitory synaptic conductance |
| E_L | -87.2 mV | Leak reversal potential |
| $\beta_{E_L}$ | 0.05 | Magnitude of dopamine effect on resting potential |
| E_ex | 0 mV | Excitatory reversal potential |
| E_in | -64 mV | Inhibitory reversal potential |
| I_e | 128 pA | Constant input current |
| C_m | 192 pF (Gertler et al., 2008) | Membrane capacitance |
| g_L | 8.04 nS (Gertler et al., 2008) | Leak conductance |
| $\beta_{V_{th}}$ | 0.205 | Magnitude of dopamine effect on threshold potential |
| t_ref | 2 ms | Duration of refractory period |

**Table 4:**
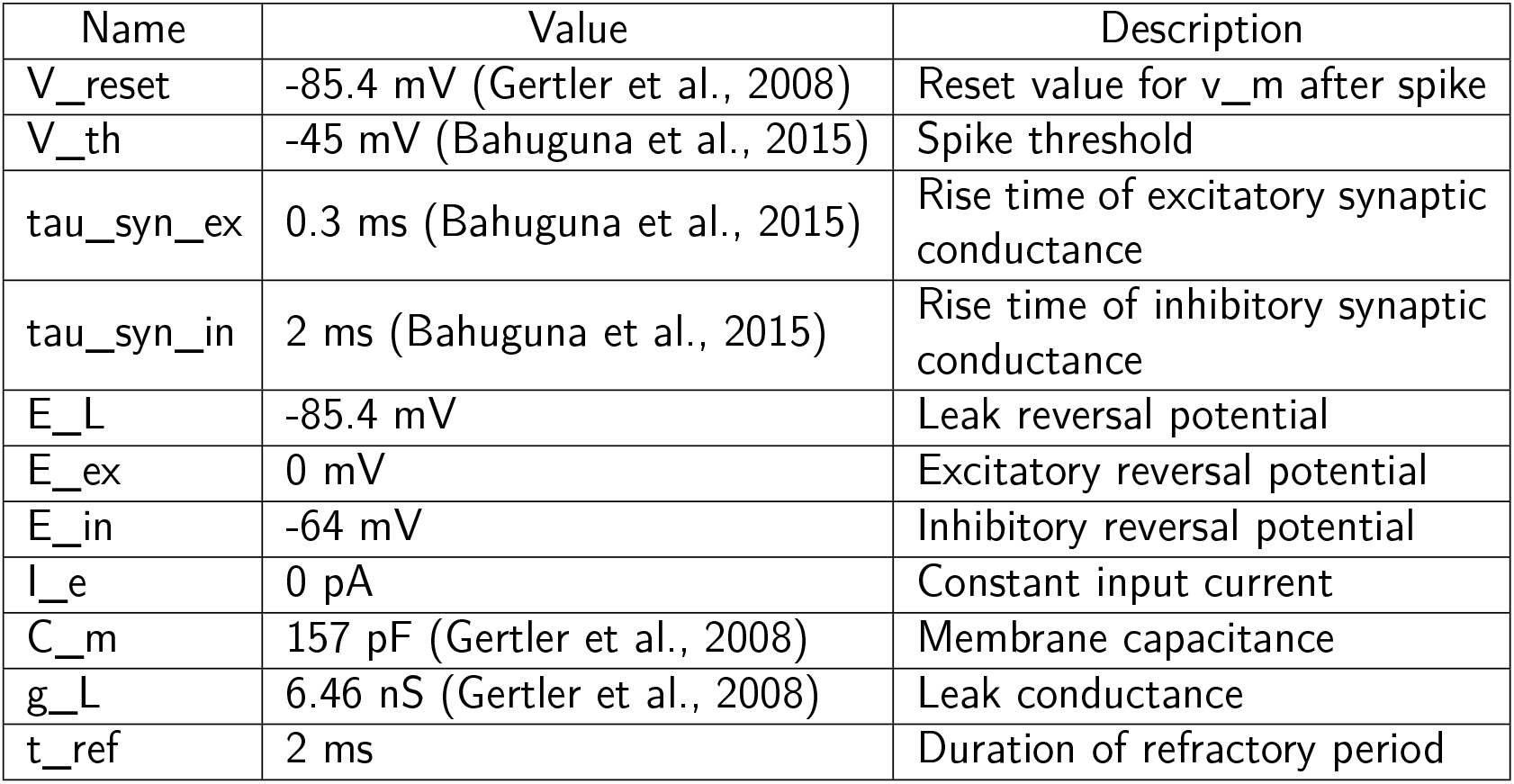
D2-SPN neuron parameters (leaky integrate and fire model).

**Table 5:** FSI neuron parameters (leaky integrate and fire model).

| Name | Value | Description |
| --- | --- | --- |
| V_reset | -65 mV (Klaus et al., 2011) | Reset value for v_m after spike |
| V_th | -54 mV (Bahuguna et al., 2015) | Spike threshold |
| tau_syn_ex | 0.3 ms (Bahuguna et al., 2015) | Rise time of excitatory synaptic conductance |
| tau_syn_in | 2 ms (Bahuguna et al., 2015) | Rise time of inhibitory synaptic conductance |
| E_L | -65 mV | Leak reversal potential |
| E_ex | 0 mV | Excitatory reversal potential |
| E_in | -76 mV | Inhibitory reversal potential |
| I_e | 0 pA | Constant input current |
| C_m | 700 pF (Klaus et al., 2011) | Membrane capacitance |
| g_L | 16.67 nS (Russo et al., 2013) | Leak conductance |
| $\beta_{E_L}$ | -0.078 (Lindahl and Kotaleski, 2016) | Magnitude of dopamine effect on resting potential |
| t_ref | 2 ms | Duration of refractory period |

**Table 8:** STN neuron parameters (leaky integrate and fire model).

| Name | Value | Description |
| --- | --- | --- |
| V_reset | -70 mV (Lindahl and Kotaleski, 2016) | Reset value for v_m after spike |
| V_th | -64 mV (Lindahl and Kotaleski, 2016) | Spike threshold |
| tau_syn_ex | 0.33 ms | Rise time of excitatory synaptic conductance |
| tau_syn_in | 1.5 ms | Rise time of inhibitory synaptic conductance |
| E_L | -80.2 mV (Lindahl and Kotaleski, 2016) | Leak reversal potential |
| E_ex | -10 mV | Excitatory reversal potential |
| E_in | -84 mV | Inhibitory reversal potential |
| I_e | 1 pA | Constant input current |
| C_m | 60 pF (Lindahl and Kotaleski, 2016) | Membrane capacitance |
| g_L | 10 nS (Lindahl and Kotaleski, 2016) | Leak conductance |
| t_ref | 2 ms | Duration of refractory period |

GPe-TA, GPe-TI and SNr neurons were modelled as a LIF neuron with exponential adaptation (AdEx). The subthreshold dynamics of these neurons was defined as:

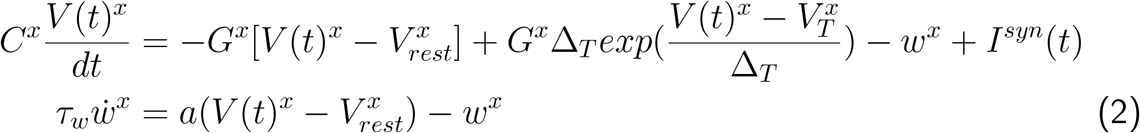

where *x* ∈ {GPe-TA, GPe-TI, SNr}. In equation 2, 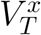 represents the spike-threshold, and *a* and *b* denote subthreshold and spike-triggered adaptation, respectively. Given equation 2, when *V*^*x*^ reaches the spike-cutoff potential then a spike is generated and *V*^*x*^, as well as *w*^*x*^ are reset at values *V*_*rest*_, *w*^*x*^ + *b*, respectively. *I*^*syn*^(*t*) models the total synaptic input current received by the neuron (see Figure 1 for the various sources of inputs to these neurons).

The neural parameters for GPe-TA, GPe-TI and SNr neurons are given in the Table 6, 7, and 9, respectively.

**Table 6:** GPe-TA neuron parameters (Lindahl and Kotaleski, 2016) (adaptive exponential integrate and fire model).

| Name | Value | Description |
| --- | --- | --- |
| a | 2.5 nS | Subthresholded adaption |
| b | 105 pA | Spike triggered adaption |
| $\beta_{E_L}$ | -0.181 | Magnitude of dopamine effect on resting potential |
| $\Delta_T$ | 2.55 ms | Slope factor |
| tau_w | 20 ms | Adaption time costant |
| V_reset | -60 mV | Reset value for v_m after spike |
| V_th | -54.7 mV | Spike initiation threshold |
| tau_syn_ex | 1 ms | Rise time of excitatory synaptic conductance |
| tau_syn_in | 5.5 ms | Rise time of inhibitory synaptic conductance |
| E_L | -55.1 mV | Leak reversal potential |
| E_ex | 0 mV | Excitatory reversal potential |
| E_in | -65 mV | Inhibitory reversal potential |
| I_e | 1 pA | Constant input current |
| C_m | 60 pF | Membrane capacitance |
| g_L | 1 nS | Leak conductance |
| t_ref | 2 ms | Duration of refractory period |

**Table 7:** GPe-TI neuron parameters (Lindahl and Kotaleski, 2016) (adaptive exponential integrate and fire model).

| Name | Value | Description |
| --- | --- | --- |
| a | 2.5 nS | Subthresholded adaption |
| b | 70 pA | Spike triggered adaption |
| $\beta_{E_L}$ | -0.181 | Magnitude of dopamine effect on resting potential |
| $\Delta_T$ | 1.7 ms | Slope factor |
| tau_w | 20 ms | Adaption time costant |
| V_reset | -60 mV | Reset value for v_m after spike |
| V_th | -54.7 mV | Spike initiation threshold |
| tau_syn_ex | 4.8 ms | Rise time of excitatory synaptic conductance |
| tau_syn_in | 1 ms | Rise time of inhibitory synaptic conductance |
| E_L | -55.1 mV | Leak reversal potential |
| E_ex | 0 mV | Excitatory reversal potential |
| E_in | -65 mV | Inhibitory reversal potential |
| I_e | 12 pA | Constant external input current |
| C_m | 40 pF | Membrane capacitance |
| g_L | 1 nS | Leak conductance |
| t_ref | 2 ms | Duration of refractory period |

**Table 9:** SNr neuron parameters (adaptive exponential integrate and fire model).

| Name | Value | Description |
| --- | --- | --- |
| a | 3 nS (Lindahl and Kotaleski, 2016) | Subthresholded adaption |
| b | 200 pA (Lindahl and Kotaleski, 2016) | Spike triggered adaption |
| $\beta_{EL}$ | -0.0896 (Lindahl and Kotaleski, 2016) | Magnitude of dopamine effect on resting potential |
| $\Delta_T$ | 1.6 ms | Slope factor |
| tau_w | 20 ms (Lindahl and Kotaleski, 2016) | Adaption time costant |
| V_reset | -65 mV (Lindahl and Kotaleski, 2016) | Reset value for v_m after spike |
| V_th | -55.2 mV (Lindahl and Kotaleski, 2016) | Spike initiation threshold |
| tau_syn_ex | 5.7 ms | Rise time of excitatory synaptic conductance |
| tau_syn_in | 2.04 ms | Rise time of inhibitory synaptic conductance |
| E_L | -55.8 mV (Lindahl and Kotaleski, 2016) | Leak reversal potential |
| E_ex | 0 mV | Excitatory reversal potential |
| E_in | -80 mV | Inhibitory reversal potential |
| I_e | 0 mV | Constant external input current |
| C_m | 80 pF (Lindahl and Kotaleski, 2016) | Membrane capacitance |
| g_L | 3 nS (Lindahl and Kotaleski, 2016) | Leak conductance |
| t_ref | 2 ms | Duration of refractory period |

### Synapse model

Neurons were connected using static conductance-based synapses. Each incoming spike elicited an alpha function shaped conductance transient, after a fixed delay since the spike in the pre-synaptic neurons. The time course of the conductance transient was given as:

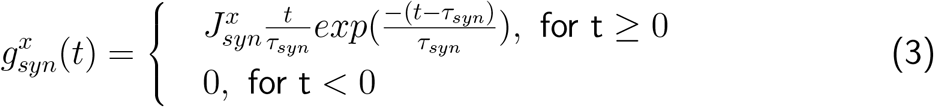

where *syn* ∈ {exc, inh} and *x* ∈ *D*1 − *SPN, D*2 − *SPN, FSI, GPe* − *TA, GPe* − *TI, STN, SNr*. In equation 3, 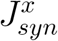 is the peak of the conductance transient and 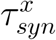 is synaptic time constant. Each incoming synaptic current induces current transient as given by:

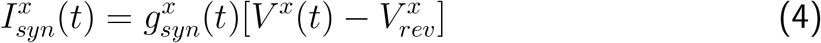

Where 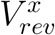 is the reversal potential of the synapse for a neuron in population *x* ∈ {*D*1 − *SPN, D*2 − *SPN, FSI, GPe* − *TA, GPe* − *TI, STN, SNr*}. All synaptic parameters are specified in Table 2.

**Table 2:** Synaptic weight and delay parameters in healthy condition.

| Weight | Values (nS) | Delay | Values (ms) |
| --- | --- | --- | --- |
| $g_{D1-SPN}^{D1-SPN}$ | -0.15 (Lindahl and Kotaleski, 2016) | $\Delta_{D1-SPN}^{D1-SPN}$ | 1.7 |
| $g_{D2-SPN}^{D1-SPN}$ | -0.375 (Lindahl and Kotaleski, 2016) | $\Delta_{D2-SPN}^{D1-SPN}$ | 1.7 |
| $g_{D1-SPN}^{D2-SPN}$ | -0.45 (Lindahl and Kotaleski, 2016) | $\Delta_{D1-SPN}^{D2-SPN}$ | 1.7 |
| $g_{D2-SPN}^{D2-SPN}$ | -0.35 (Lindahl and Kotaleski, 2016) | $\Delta_{D2-SPN}^{D2-SPN}$ | 1.7 |
| $g_{D1-SPN}^{FSI}$ | -2.6 (Bahuguna et al., 2015) | $\Delta_{D1-SPN}^{FSI}$ | 1.7 |
| $g_{D2-SPN}^{FSI}$ | -2.6 (Bahuguna et al., 2015) | $\Delta_{D2-SPN}^{FSI}$ | 1.7 |
| $g_{D1-SPN}^{GPe-TA}$ | -0.02 | $\Delta_{D1-SPN}^{GPe-TA}$ | 7 |
| $g_{D2-SPN}^{GPe-TA}$ | -0.04 | $\Delta_{D2-SPN}^{GPe-TA}$ | 7 |
| $g_{FSI}^{FSI}$ | -0.4 | $\Delta_{FSI}^{FSI}$ | 1.7 |
| $g_{FSI}^{GPe-TA}$ | -0.25 | $\Delta_{FSI}^{GPe-TA}$ | 7 |
| $g_{FSI}^{GPe-TI}$ | -1 | $\Delta_{FSI}^{GPe-TI}$ | 7 |
| $g_{SNr}^{GPe-TI}$ | -52.5 | $\Delta_{SNr}^{GPe-TI}$ | 3 |
| $g_{SNr}^{D1-SPN}$ | -15 | $\Delta_{SNr}^{D1-SPN}$ | 7 |
| $g_{SNr}^{STN}$ | 4.78 | $\Delta_{SNr}^{STN}$ | 4 |
| $g_{GPe-TI}^{D2-SPN}$ | -1.08 | $\Delta_{GPe-TI}^{D2-SPN}$ | 7 |
| $g_{GPe-TA}^{STN}$ | 0.24 | $\Delta_{GPe-TA}^{STN}$ | 2 |
| $g_{GPe-TI}^{STN}$ | 0.175 | $\Delta_{GPe-TI}^{STN}$ | 2 |
| $g_{GPe-TA}^{GPe-TA}$ | -0.11 | $\Delta_{GPe-TA}^{GPe-TA}$ | 1 |
| $g_{GPe-TI}^{GPe-TA}$ | -1.3 | $\Delta_{GPe-TI}^{GPe-TA}$ | 1 |
| $g_{GPe-TA}^{GPe-TI}$ | -0.35 | $\Delta_{GPe-TA}^{GPe-TI}$ | 1 |
| $g_{GPe-TI}^{GPe-TI}$ | -1.3 | $\Delta_{GPe-TI}^{GPe-TI}$ | 1 |
| $g_{STN}^{GPe-TI}$ | -0.3 | $\Delta_{STN}^{GPe-TI}$ | 1 |

### Basal ganglia network

The basal ganglia (BG) comprises of striatum, subthalamic nucleus (STN), globus pallidus externa (GPe), substantia nigra pars reticulata (SNr) and globus pallidus interna (GPi) in primates or entopeduncular nucleus (EPN) in rodents (Figure 1). To model BG, we adapted a previously published model by Lindahl and Kotaleski (2016). However unlike that model (Lindahl and Kotaleski, 2016), here we reduced the time complexity of our proposed network by scaling down the size of striatum (D1-SPN, D2-SPN, FSI). Also a few synaptic and neural parameters were adjusted to achieve the network performance in healthy and PD conditions. The main differences between these two models are detailed in the later part of methods section.

Our reduced model of the BG consisted of 6539 neurons. Number of neurons in each sub-population, number of connections and synaptic connectivity parameters are provided in Table 1.

**Table 1:** Network and connection parameters (Lindahl and Kotaleski, 2016; Bahuguna et al., 2015).

| Name | Value | Description |
| --- | --- | --- |
| $N_{network}$ | 6539 | Network size |
| $N_{network}^{D1-SPN}$ | 2000 | Size of D1-SPN population |
| $N_{network}^{D2-SPN}$ | 2000 | Size of D2-SPN population |
| $N_{network}^{FSI}$ | 80 | Size of FSI population |
| $N_{network}^{STN}$ | 388 | Size of STN population |
| $N_{network}^{GPe-TA}$ | 329 | Size of GPe-TA population |
| $N_{network}^{GPe-TI}$ | 988 | Size of GPe-TI population |
| $N_{network}^{SNr}$ | 754 | Size of SNr population |
| $K_{D1-SPN}^{D1-SPN}$ | 364 | Number of D1-SPN connections on each D1-SPN |
| $K_{D2-SPN}^{D1-SPN}$ | 84 | Number of D1-SPN connections on each D2-SPN |
| $K_{D1-SPN}^{D2-SPN}$ | 392 | Number of D2-SPN connections on each D1-SPN |
| $K_{D2-SPN}^{D2-SPN}$ | 504 | Number of D2-SPN connections on each D2-SPN |
| $K_{D1-SPN}^{FSI}$ | 16 | Number of FSI connections on each D1-SPN neuron |
| $K_{D2-SPN}^{FSI}$ | 11 | Number of FSI connections on each D2-SPN neuron |
| $K_{D1-SPN}^{GPe-TA}$ | 10 | Number of GPe-TA connections on each D1-SPN neuron |
| $K_{D2-SPN}^{GPe-TA}$ | 10 | Number of GPe-TA connections on each D2-SPN neuron |
| $K_{FSI}^{FSI}$ | 10 | Number of FSI connections on each FSI neuron |
| $K_{FSI}^{GPe-TA}$ | 10 | Number of GPe-TA connections on each FSI neuron |
| $K_{FSI}^{GPe-TI}$ | 10 | Number of GPe-TI connections on each FSI neuron |
| $K_{SNr}^{GPe-TI}$ | 32 | Number of GPe connections on each SNr neuron |
| $K_{SNr}^{D1-SPN}$ | 500 | Number of D1-SPN connections on each SNr neuron |
| $K_{SNr}^{STN}$ | 30 | Number of STN connections on each SNr neuron |
| $K_{GPe-TI}^{D2-SPN}$ | 500 | Number of D2-SPN connections on each GPe-TI neuron |
| $K_{GPe-TA}^{STN}$ | 30 | Number of STN connections on each GPe-TA neuron |
| $K_{GPe-TI}^{STN}$ | 30 | Number of STN connections on each GPe-TI neuron |
| $K_{GPe-TA}^{GPe-TA}$ | 5 | Number of GPe-TA reciprocal connections |
| $K_{GPe-TI}^{GPe-TA}$ | 5 | Number of GPe-TA connections on each GPe-TI neuron |
| $K_{GPe-TA}^{GPe-TI}$ | 25 | Number of GPe-TI connections on each GPe-TA neuron |
| $K_{GPe-TI}^{GPe-TI}$ | 25 | Number of GPe-TI reciprocal connections |
| $K_{STN}^{GPe-TI}$ | 30 | Number of GPe-TI connections on each STN neuron |

### Dopamine induced changes in neuron and synapse parameters

To model the effect of dopamine we followed the approach taken by Lindahl and Kotaleski (2016). Dopaminergic effects on SPNs, FSIs, STN, GPe and SNr neurons and their synaptic connections were modelled by modulating parameters such as the resting state potentials (*E*_*L*_), spike threshold (*V*_*th*_), and synaptic strengths. To model dopamine modulation we varied the parameter *α*_*dop*_ from 0 to 1. We set *α*_*normal*_ = 0.8 for normal and *α*_*dop*_ = 0 for PD conditions.

#### Dopamine effects on neuron properties

In D1-SPNs, D1 type dopamine receptor activation not only shows a hyperpolarizing effect by increasing potassium inward rectifier (KIR) current, but also induces depolarizing effects on the resting membrane potential (Gruber et al., 2003). We modelled these two contributions by changing the spike threshold and resting membrane potential:

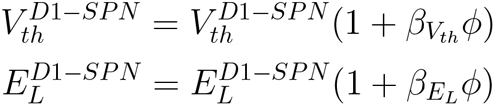

where *ϕ* = *α*_*dop*_ *α*_*normal*_. The parameters 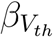 and 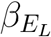 (see table 3) were chosen based on Humphries et al. (2009). Although Planert et al. (2013) suggested that dopamine concentration modulates the excitability of D2-SPN, in low-dopamine state with 60*µM* dopamine concentration, no significant changes in their excitability was observed. Therefore, following the reasoning given by Lindahl and Kotaleski (2016) in this model we also ignored the effects of dopamine on the D2-SPNs. We modelled the dopaminergic depolarizing effect induced through D1 type receptor activation on the FSIs, by modulating their resting membrane potential:

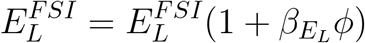

where 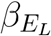 (see Table 5) was set such that 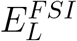 at low dopamine level was 5 mV lower than that of the high dopamine level (Bracci et al., 2002).

The dopaminergic depolarizing effects on the GPe neurons (both TA and TI) are manifested as up-regulation of the hyperpolarization-activated cyclic nucleotide–gated (HCN) channels (Chan et al., 2011) which essentially results in a change in the resting membrane potential of the neurons. To mimic this effect we changed the resting membrane potential of the GPe neurons in the following manner:

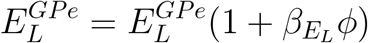

The values of 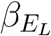 for both the GPe-TA and GPe-TI neurons (see Tables 6 and 7) were set such that the resting state potential of the GPe neurons at low dopamine level was 10 mV lower than that of its value at high dopamine level.

Dopaminergic effect on the SNr neurons (Zhou et al., 2009) was realized by changing their resting membrane potential:

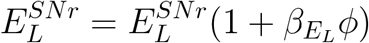

Where 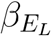 (see table 9) was taken such that the resting potential at low dopamine level was 5 mV lower than its value at high dopamine level.

The scaling factors *β*_*i*_ (*i* ∈ {*E*_*L*_, *V*_*th*_}), for the linear modulation (*ϕ* = *α*_*dop*_ − *α*_*normal*_) were tuned for each parameter to match their experimentally reported results in both normal and PD conditions (rodent models).

#### Dopamine effects on synaptic weights

High dopamine strengthens cortical projection on to D1-SPN and weakens cortical projections on to D2-SPN (Hernández-Echeagaray et al., 2004). The decrease in connectivity both in terms of synaptic strength and number of recurrent connections among SPNs is also attributed to dopamine depletion (Taverna et al., 2008). In addition, dopamine is reported to reduce the strength of GABAergic synapses (Bracci et al., 2002) between FSI-FSI and dopamine depletion increases the number of connections between FSI and D2-SPN (Gittis et al., 2011), but not D1-SPN. Within the GPe, dopamine depletion strengthens the GPe↔GPe (Miguelez et al., 2012) and GPe→FSI connections, through the activation of D2 receptors. In addition to that it also strengthens the GPe-TA→SPN synapses (Glajch et al., 2016). Dopamine depletion also strengthens the D2-SPN projections on to GPe neurons through reduced D2-receptor activation (Chuhma et al., 2011). Similarly, high dopamine reduces the strength of STN→GPe synapses (Hernández et al., 2006). Dopamine is also responsible for reducing the synaptic efficacy in GPe-TI→STN synapses (Baufreton and Bevan, 2008). In addition, high dopamine also weakens cortical synapses on to STN neurons (Shen and Johnson, 2006).

On the other hand, at high dopamine, the D1-SPN to SNr connection was facilitated, hence the *I*_*GABA*_ from D1-SPN to SNr was modelled to reflect the same (Chuhma et al., 2011).

Dopaminergic effect on the synaptic strength 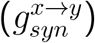 was modelled as 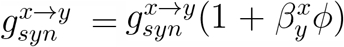where *x, y* ∈ {FSI, D1-SPN, D2-SPN, STN, Cortex, GPe, SNr} and the values of 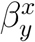 were given in the Table 10.

**Table 10:** Synaptic dopamine parameters. To obtain a triphasic response in PD condition we had to change a few parameters of the network tuned in default PD state (biphasic). These changes are marked in boldface.

| Name | Value in PD-Biphasic | Value in PD-Triphasic |
| --- | --- | --- |
| $\beta_{FSI}^{FSI}$ | -1.27 (Lindahl and Kotaleski, 2016) | -1.27 |
| $\beta_{FSI}^{GPe}$ | -0.53 (Lindahl and Kotaleski, 2016) | -0.53 |
| $\beta_{GPe}^{GPe}$ | -0.83 (Lindahl and Kotaleski, 2016) | -0.83 |
| $\beta_{GPe-TI}^{D2-SPN}$ | -1.00 | **-0.48** |
| $\beta_{GPe}^{STN}$ | -0.3 | -0.3 |
| $\beta_{D1-SPN}^{Cortex}$ | 1.04 (Lindahl and Kotaleski, 2016) | 1.04 |
| $\beta_{D2-SPN}^{Cortex}$ | -0.26 (Lindahl and Kotaleski, 2016) | -0.26 |
| $\beta_{D2-SPN}^{FSI}$ | -0.90 (Lindahl and Kotaleski, 2016) | -0.90 |
| $\beta_{SPN}^{SPN}$ | 0.88 (Lindahl and Kotaleski, 2016) | 0.88 |
| $\beta_{D1-SPN}^{GPe-TA}$ | -1.22 (Lindahl and Kotaleski, 2016) | -1.22 |
| $\beta_{D2-SPN}^{GPe-TA}$ | -1.15 (Lindahl and Kotaleski, 2016) | -1.15 |
| $\beta_{SNr}^{D1-SPN}$ | 0.42 | **0.56** (Lindahl and Kotaleski, 2016) |
| $\beta_{STN}^{Cortex}$ | -1.15 | -1.15 |
| $\beta_{STN}^{GPe}$ | -0.54 | **-0.24** (Lindahl and Kotaleski, 2016) |

### External inputs

In our network model, all the neuronal populations received uncorrelated excitatory Poisson input spike-train, mimicking background inputs either from cortex or from thalamus. The input rates were tuned both in normal and PD conditions to ensure that the basal firing rates of different subnuclei were consistent with the *in vivo* recordings in rats, e.g., in normal condition baseline firing rate (in Hz) of D1-SPN and D2-SPN ∈ [0.01, 2.0] (Miller et al., 2008; Lindahl and Kotaleski, 2016), FSI ∈ [10, 20] (Gage et al., 2010), STN ∈ [10, 13] (Fujimoto and Kita, 1993; Paz et al., 2005) and SNr ∈ [20, 35] (Kita and Kita, 2011; Benhamou and Cohen, 2014). The baseline activities of GPe-TA and GPe-TI are matched to the work done by Mallet et al. (2008). Similarly, in PD condition, frequencies of the background noise (in Hz) were also tuned to achieve range of basal firing rate of D1-SPN ∈ [0.1, 0.5], D2-SPN ∈ [1, 2], GPe-TA ∈ [12, 16] (de la Crompe et al., 2020), GPe-TI ∈ [17, 20] (de la Crompe et al., 2020) and STN ∈ [26, 29] (de la Crompe et al., 2020). For SNr, Sano and Nambu (2019) claimed a decrease of basal firing rate in PD conditions, however others (Kita and Kita, 2011; Ruskin et al., 2002) had not observed firing rate changes in PD state. Given this, we kept basal firing rate of SNr same, as it is in normal state ∈ [29, 32].

To characterize the effect of a transient cortical stimulation on the neuronal responses of the SNr, we stimulated striatal and STN neurons with a brief stimulus pulse which amounted to injection of one spike to all the stimulated neurons. The cortical stimulation was modelled by injecting a single spike in D1-SPN, D2-SPN, FSI and STN neurons at time *T*_*stimulation*_ (the stimulation time). The spike was injected in a different fraction of neurons. This input was modelled by using the spike_generator device in NEST (Gewaltig and Diesmann, 2007). Because the transient stimulation was modelled as injection of spike, we could control the strength of input stimulation by varying the amplitude of the excitatory postsynaptic potential (EPSP) generated by the injected spike. Moreover, this allowed us to modulate the strength of input in relation to dopamine levels (see the subsection **Dopamine effects on synaptic weights** for how dopamine affected synaptic weights). Transient response was measured in both normal and PD conditions.

### Main differences between our model and the one proposed by Lindahl and Kotaleski (2016)

Here we used the model by Lindahl and Kotaleski (2016); however, we made a few changes in the neuron and synapse models and changed the number of neurons in some of the BG subnetworks. Unlike their model (Lindahl and Kotaleski, 2016), striatal and STN neurons were modelled as simple LIF neurons without any kind of adaptation and, all the synapses were static as opposed to the dynamic ones. In addition, we also reduced the size of striatal neuronal population. To compensate for this change, we changed the synaptic strengths and a few neuronal model parameters, such that the average synaptic input to GPe and GPi/SNr neurons was identical to the model used by Lindahl and Kotaleski (2016). This ensured that the model was operating in the same regime as that of the model by Lindahl and Kotaleski (2016). In addition, we assumed that all the synapses are static. Finally, to generate triphasic shaped transient responses, we also changed the values of 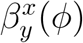 (see Table 10). Besides these changes, we followed the model closely while modelling the effects of dopamine on neuron and synapse parameters.

### Limitations of the model

Unlike in the experimental data, in our model all neurons responded with similar response profile. This is because the model is homogeneous in terms of neuron and synapse properties. It was important to keep the model homogeneous in order to isolate the various interactions that lead to triphasic or other shapes of transient response. Furthermore, all synapses are static in this model. We made this choice to reduce the computational demands of the simulations. We note that Lindahl et al. (2013) suggested that synaptic short term plasticity is important for the triphasic response when STN is stimulated. However, as we show in this study, triphasic responses do not rely on synaptic short-term dynamics. Moreover, to the best of our knowledge, there is no experimental evidence for short-term plasticity to be the cause of triphasic response. We have only considered effects of changes in the dopamine baseline. Transient responses of BG could also be accompanied by phasic change in DA levels. Such effects have been ignored. In addition, we have ignored the effect of low-dopamine on D2-SPNs, although we mimicked this effect indirectly by increasing their basline activity in PD condition. We have only modelled the fast spiking interneurons and ignored other types of interneurons. Only recently a detailed microcircuit has been modelled with numerical simulations (Hjorth et al., 2020). In future studies it may be possible to use a reduced version of that network for BG modelling. Finally, our model does not address the changes in the spatio-temporal dynamics of BG nuclei given cortical stimulation, as connectivity within each subnetwork is independent of spatial distances among the neurons.

### Simulation tools

All the simulations were performed using the simulator NEST (Jordan et al., 2019). All differential equations were integrated using Runga-Kutta method with a time step of 0.1 ms. Simulation code will be made available on github upon publication of the manuscript.

### Data Analysis

#### Transient response analysis

To get better estimate of the transient response we performed 100 trials and recorded the response over 1200 ms. The timing of the transient input was randomly chosen between 700 ms and 900 ms for every trial, which was later time aligned at the stimulation point for further analysis. Note that, the stimulation point was chosen between 700 ms to 900 ms to discard the initial transient noisy effect which was mainly caused by the instability of the network from our analysis. To understand effect of the transient stimulation on the SNr activities, the neuronal responses of SNr neurons were observed before and after the cortical stimulation point *T*_*stimulation*_. As mentioned earlier *T*_*stimulation*_ was randomly chosen for every trial. A 350 ms window size was defined around *T*_*stimulation*_ to extract responses from each trial. For this, we used a time window of 100 ms before and 250 ms after the stimulation point.

The responses were evaluated by constructing peristimulus time histograms (PSTH), using 1 ms rectangular bins for each trial data. To analyze the transient response of the SNr, both in normal and PD conditions, the PSTH were divided into four zones (see Figure 2). These zones consisted of two excitatory (EE and LE for early and late, respectively) and two inhibitory (EI and LI) zones. The change in firing activity was marked as excitation or inhibition, if the firing rate was significantly different from the baseline (*P* < 0.05, one-tailed Z-test) for at least two consecutive time bins (2 ms) (Sano et al., 2013). The latency of each zone was measured as the time when the first bin exceeded the baseline. Similarly, the zone terminated when activities during two consecutive bins fell below the significance level. The end time was determined as the time of the last bin exceeding the significance level. The total time duration from the first bin to the last bin (of a significant response) was considered as the duration of each zone. The sum of heights of bins within a particular zone is considered as the area as well as the strength of the zone, whereas the area per unit time (area/time) indicates the average strength of that zone. Thus, we extracted the following features from PSTH for each zone: latency (*L*), duration (*D*), absolute area indicating strength (*A*) of that zone, mean (*H*_*µ*_), and standard deviation (*H*_*σ*_) of bin-heights. In addition, we also measured the peak amplitudes (*H*_*p*_) of each zone (i.e. *H*_*p*_ ∈ {*H*_*EE*−*max*_, *H*_*LE*−*max*_, *H*_*EI*−*min*_, *H*_*LI*−*min*_}).

**Figure 2:**
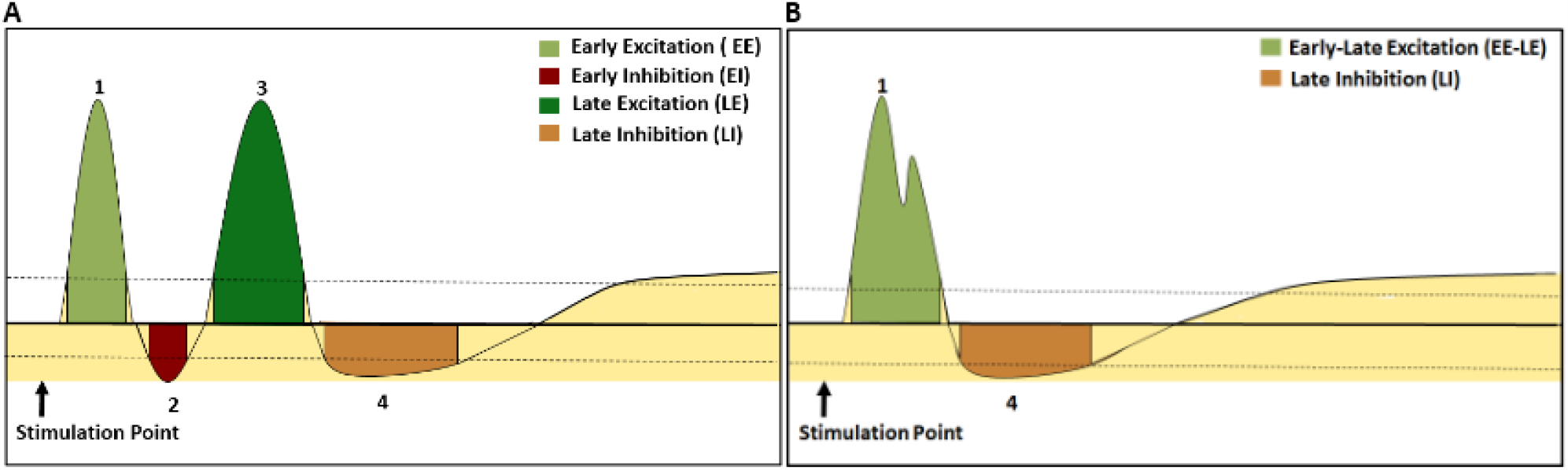
Characterization of transient responses. (**A**) A schematic representation of cortical stimulation induced triphasic response patterns in the SNr (see in healthy state). The triphasic response consists of early excitation (EE), early inhibition (EI), late excitation (LE) and a late inhibition (LI). (**B**) A schematic representation of biphasic shaped transient response patterns in the SNr (corresponding to what is seen in PD condition). It consists of EE, LE, and a LI. The horizontal bold line and two dotted lines denote the prestimulus mean (basal) firing rate and 95% confidence interval, respectively

Finally, *F*_*i*_ (i ∈ EE, EI, LE, LI})) is a six dimensional vector ({*L, D, A, H*_*µ*_, *H*_*σ*_, *H*_*p*_}). *F*_*i*_s were computed for each zone in different network conditions. The similarity between a network tuned in healthy condition (TestNetwork) and PD condition was calculated using Euclidean distance metric:

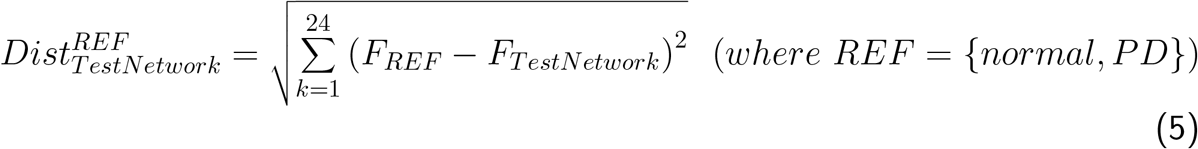

In order to observe the statistical variation of the above features within the normal and PD conditions, a sub-population of SNr neurons were considered. From the whole population of SNr neurons, a percentage of neurons (*N*_*S*_%) was randomly chosen to represent an observation. The responses of these sub-population of SNr neurons were averaged over multiple trials (*O*_*S*_ = 100 in the present simulation) and then the features of the four zones were extracted. The choice of the sub-population of *N*_*S*_% was also varied for every observations and a large number of such observations (*O*_*S*_) were made. These observations were used to derive the mean and standard deviation of the features for each of the above mentioned EE, EI, LE and LI zones. Here, we have considered *O*_*S*_ = 100 and *N*_*S*_ = 50%.

In experimental studies (Ozaki et al., 2017; Sano and Nambu, 2019), in healthy state, the transient response is often chracterized by dividing the response in three zones EE, EI and LE. In low-dopamine state the experimentally observed biphasic response pattern could consist of any two zones out of the three zones (EE, EI and LE) (Sano and Nambu, 2019). Experimental data also shows that in both normal and PD conditions, LE is followed by an LI zone Kita and Kita (2011). Therefore, here we defined four zones to characterize the transient response in healthy state (Figure 2 A). So even though we have defined four zones, we still refer to it as a triphasic response in order to be consistent with the terminology used in the literature. In our simulations, biphasic response (Figure 2 B) observed in PD condition consisted of EE and LE, while the EI zone was not observed. We merged the EE and LE zones together for computing the features of the PD-biphasic response.

### Global network activity

The oscillatory behaviour of population activities was assessed in PD as well as normal conditions. We ran the simulation for longer duration (5*seconds*), without any transient input, to allow the oscillations to set in to their steady state. This experiment was also carried out over 100 trials.

Synchrony in the firing rates of a neuronal population was estimated using Fano Factor (*FF*_*pop*_) (Kumar et al., 2008):

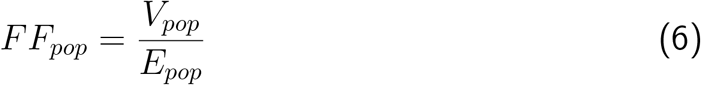

where *E*_*pop*_ and *V*_*pop*_ are the mean and variance of the population activity, respectively. For an uncorrelated ensemble of Poisson processes, *FF*_*pop*_ = 1 and when neurons tend to correlate, *FF*_*pop*_ > 1. Here, we binned the neuronal activity using rectangular bins of 3 ms duration. This window size was similar to the one used in previous studies (Mallet et al., 2008; Lindahl and Kotaleski, 2016).

To determine the strength of oscillatory neuronal activities in the *β*-band we estimated the oscillation index (*OI*_*pop*_). To this end, we estimated the spectrum of the population activity (*S*_*pop*_(*f*)). As we used 3 ms bins to calculate the PSTH, the sampling frequency (*F*_*s*_) was 333.3 Hz. To estimate the oscillation index we measured the relative power confined in the *β*-band:

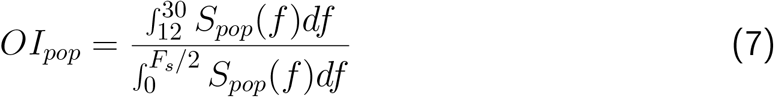

The phase relation of the firing pattern between the two types (Type-A and Type-I) of GPe nuclei as well as with STN were computed from the PSTH, having bin size of 1 ms. As we were interested in analyzing the pathological *β*-oscillation, the individual PSTH responses were bandpass filtered between 12 and 30 Hz.

Initially, at every time instance, corresponding to each bin, the instantaneous phase was calculated using the Hilbert Transform. Then the differences of the instantaneous phases were obtained between a pair of nuclei, for every 1 ms. Finally, the histogram of the difference in the phase was obtained with 100 bins in the range of 0 to *π*.

## Results

The standard feedforward model of the BG predicts that transient cortical stimulation will result in a triphasic response in the SNr as the stimulus induced activity is propagated over the direct, indirect and hyper-direct pathways. Indeed, many neurons, at least in a healthy state, do show a triphasic response *in vivo*. However, in both healthy and dopamine-depleted conditions, response pattern of a sizeable fraction of neurons deviates from the triphasic response shape (Sano and Nambu, 2019; Sano et al., 2013; Kita and Kita, 2011; Ozaki et al., 2017) indicating the role of recurrent interactions within and between BG nuclei. To understand how different neurons and network parameters shape the output of SNr when the striatum and STN are transiently stimulated, we used numerical simulations of the BG network with spiking neurons. In the model we systematically varied the dopamine level and studied how strength of different connections in the BG affects the shape of the transient response in both healthy and PD conditions. Here we set the dopamine level to 0.8 and 0.0 to tune the model into healthy and PD conditions, respectively (Lindahl and Kotaleski, 2016).

### Cortically evoked transient response in SNr

To study the response of BG to transient cortical stimulation we injected a single spike synchronously in 50% of the striatal and STN neurons (see Methods). Consistent with the predictions of a feedforward model of the BG and *in vivo* experimental data, in healthy state SNr neurons responded with a triphasic response consisting of early excitation (due to STN), inhibition (due to the D1-SPN projections), and late excitation (due to indirect pathway) i.e. the EE-EI-LE response (see Figure 3 A). By contrast, in PD condition, SNr neurons responded with a biphasic response (default PD condition), consisting of a prominent early excitation and late excitation (i.e. EE-LE, see Figure 3 B). Thus, the model suggests that dopamine induced changes (see Methods and 10) not only affect the steady-state of the BG network (i.e. *β*-oscillations) but also impair the transient inhibitory effect of the striatal input to the SNr because of weak D1-SPN projections as well as stronger activity along the indirect pathway.

**Figure 3:**
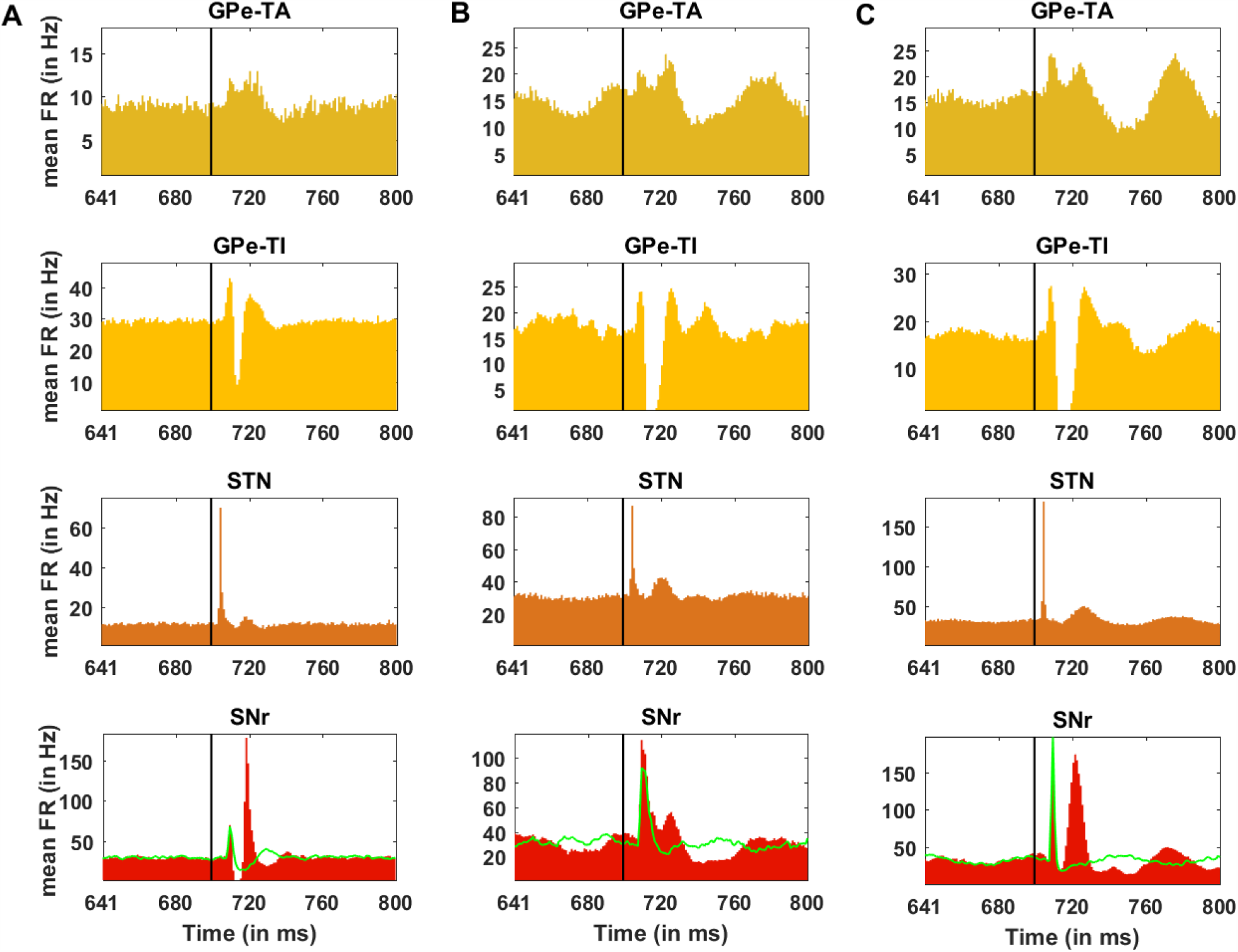
Cortically evoked responses in the GPe-TA, GPe-TI, STN and SNr. (**A**) Average PSTH (100 trials) of all neurons in GPe-TA, GPe-TI, STN and SNr. (**B**) Average PSTH (100 trials) of all neurons in GPe-TA, GPe-TI, STN and SNr in PD-biphasic condition. (**C**) Average PSTH (100 trials) of all neurons in GPe-TA, GPe-TI, STN and SNr in PD-triphasic condition. The black vertical line represents the stimulation onset. The green curve in each panel denotes the transient response of the SNr neurons in response to only STN stimulation.

Experimental data shows that even in the PD condition ≈ 40% of the SNr neurons respond in a triphasic manner (Sano and Nambu, 2019). In our model, in order to generate a triphasic response in PD condition (Figure 3 C), we needed to make additional changes than those brought in by low-dopamine. In particular, we reduced D2-SPN→GPe-TI, increased D1-SPN→SNr and reduced GPe-TI→STN connections (see Table 10 for numerical values). Note that these changes did not affect the properties of the baseline activity in a qualitative manner.

While qualitatively in both healthy and PD conditions cortical stimulation evoked an early excitation but in PD condition (both biphasic and triphasic) the duration and amplitude of early excitation were higher than that of in healthy condition. This was because dopamine depletion reduced the inhibitory effect of direct pathway and amplified the excitation of SNr neurons through the hyper-direct pathway. Moreover, in PD condition when we could generate triphasic response pattern, the duration and amplitude of early inhibition (i.e. EI) were much smaller than that of in healthy condition (early inhibition was completely absent in the biphasic responses). Finally, the late excitation phase (LE) of the triphasic response was prolonged in duration but weakened in amplitude in PD condition as compared to the healthy condition. The details of further differences in transient response properties are provided in the Table 11. The trend of the features in normal and PD conditions is consistent with the experimental data (Sano and Nambu, 2019; Ozaki et al., 2017).

**Table 11:**
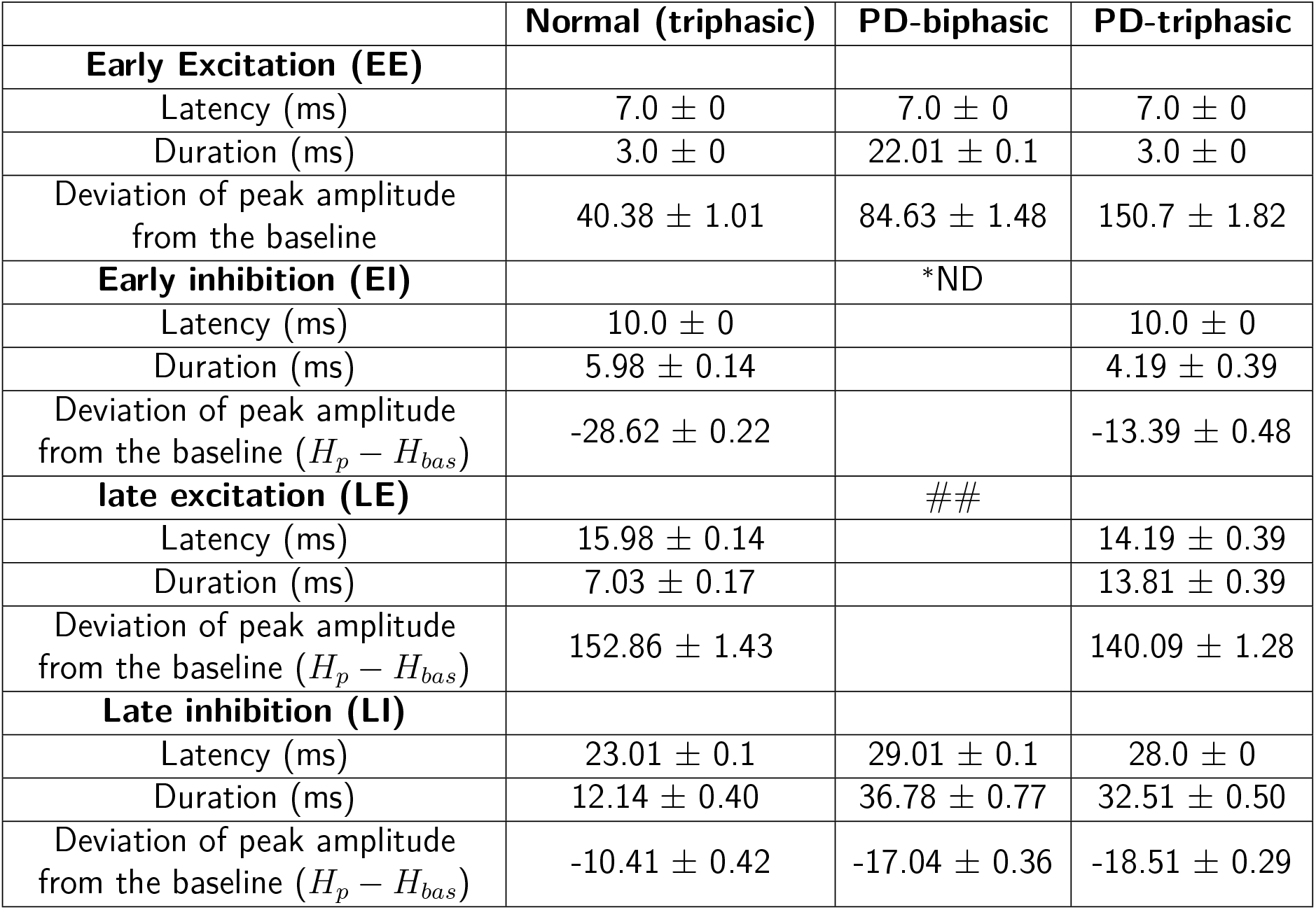
Features of the transient response of the SNr neurons. Here the variations in the features were obtained using multiple observations (100 in number) of the simulation output. In each observation, 50% of SNr neurons were randomly chosen.^##^The statistics for EE in PD-biphasic are given for the complete excitatory response comprising of both EE and LE. In this case, the EI was not detectable using statistical test, hence the two excitations (EE and LE) were merged during computation of the parameters. Here, the deviation of peak amplitude (*H*_*p*_) was measured with respect to baseline (*H*_*bas*_).^∗^ND denotes that the zone was not detected using significance test (See the subsection **Data Analysis**).

|  | Normal (triphasic) | PD-biphasic | PD-triphasic |
| --- | --- | --- | --- |
| **Early Excitation (EE)** |  |  |  |
| Latency (ms) | $7.0 \pm 0$ | $7.0 \pm 0$ | $7.0 \pm 0$ |
| Duration (ms) | $3.0 \pm 0$ | $22.01 \pm 0.1$ | $3.0 \pm 0$ |
| Deviation of peak amplitude from the baseline | $40.38 \pm 1.01$ | $84.63 \pm 1.48$ | $150.7 \pm 1.82$ |
| **Early inhibition (EI)** |  | *ND |  |
| Latency (ms) | $10.0 \pm 0$ | | $10.0 \pm 0$ |
| Duration (ms) | $5.98 \pm 0.14$ | | $4.19 \pm 0.39$ |
| Deviation of peak amplitude from the baseline ($H_p - H_{bas}$) | $-28.62 \pm 0.22$ | | $-13.39 \pm 0.48$ |
| **late excitation (LE)** |  | ## |  |
| Latency (ms) | $15.98 \pm 0.14$ | | $14.19 \pm 0.39$ |
| Duration (ms) | $7.03 \pm 0.17$ | | $13.81 \pm 0.39$ |
| Deviation of peak amplitude from the baseline ($H_p - H_{bas}$) | $152.86 \pm 1.43$ | | $140.09 \pm 1.28$ |
| **Late inhibition (LI)** |  |  |  |
| Latency (ms) | $23.01 \pm 0.1$ | $29.01 \pm 0.1$ | $28.0 \pm 0$ |
| Duration (ms) | $12.14 \pm 0.40$ | $36.78 \pm 0.77$ | $32.51 \pm 0.50$ |
| Deviation of peak amplitude from the baseline ($H_p - H_{bas}$) | $-10.41 \pm 0.42$ | $-17.04 \pm 0.36$ | $-18.51 \pm 0.29$ |

These results suggest that dopamine depletion primarily affected the early-inhibition and late-excitation zones. On one hand dopamine depletion reduced excitability of D1-SPN (Gruber et al., 2003) and reduced basal firing in D1-SPN while increasing in firing rate of D2-SPNs. Therefore, the direct pathway was weakened and resulted in reduced early inhibition (“EI”) in SNr. On the other hand the basal firing rate of GPe-TI was reduced and GPe-TA was increased (Mallet et al., 2008). This resulted in weakening of amplitude but prolonged “LE” zone.

### STN evoked transient response in the SNr

To separate the contribution of direct and hyper-direct pathways we measured the SNr response when only STN was stimulated (Figure 3 bottom row, green trace). In healthy state, consistent with experimental data (Maurice et al., 2003) and a previous modelling study (Lindahl et al., 2013), STN stimulation alone generated a triphasic response in the SNr, however there were notable differences: the EI zone was weaker, LE zone was both weaker and delayed and, LI zone was absent. Here, STN to SNr connections shaped the EE zone, GPe-TI activity gave rise to the EI zone and the LE zone was caused by STN-TI-SNr pathway. In PD condition STN stimulation induced a transient response with only EE zone (essentially, EE and LE zones observed in a healthy state were merged into a single excitatory zone). The magnitude of EE zone was much higher in the PD triphasic configuration. These results confirm that in healthy state the hyper-direct pathway shapes the EE zone, the direct pathway is responsible for the EI zone and indirect pathway shapes the LE/LI zones.

### Effect of the strength of cortical stimulation on the transient response

The aforementioned transient responses were measured by stimulating 50% of the striatal and STN population. Next, we asked whether differences in the shape of triphasic response observed in PD and healthy conditions could be reduced by stimulating more neurons. To this end, we systematically increased the number of striatal and STN neurons that received cortical stimulation (to mimic the strength of cortical stimulation). To quantify the changes in the shape of the triphasic responses we measured the duration and area per unit time (area/time) of the four zones in healthy (Figure 4 B, C) and PD condition (Figure 4 D, E).

**Figure 4:**
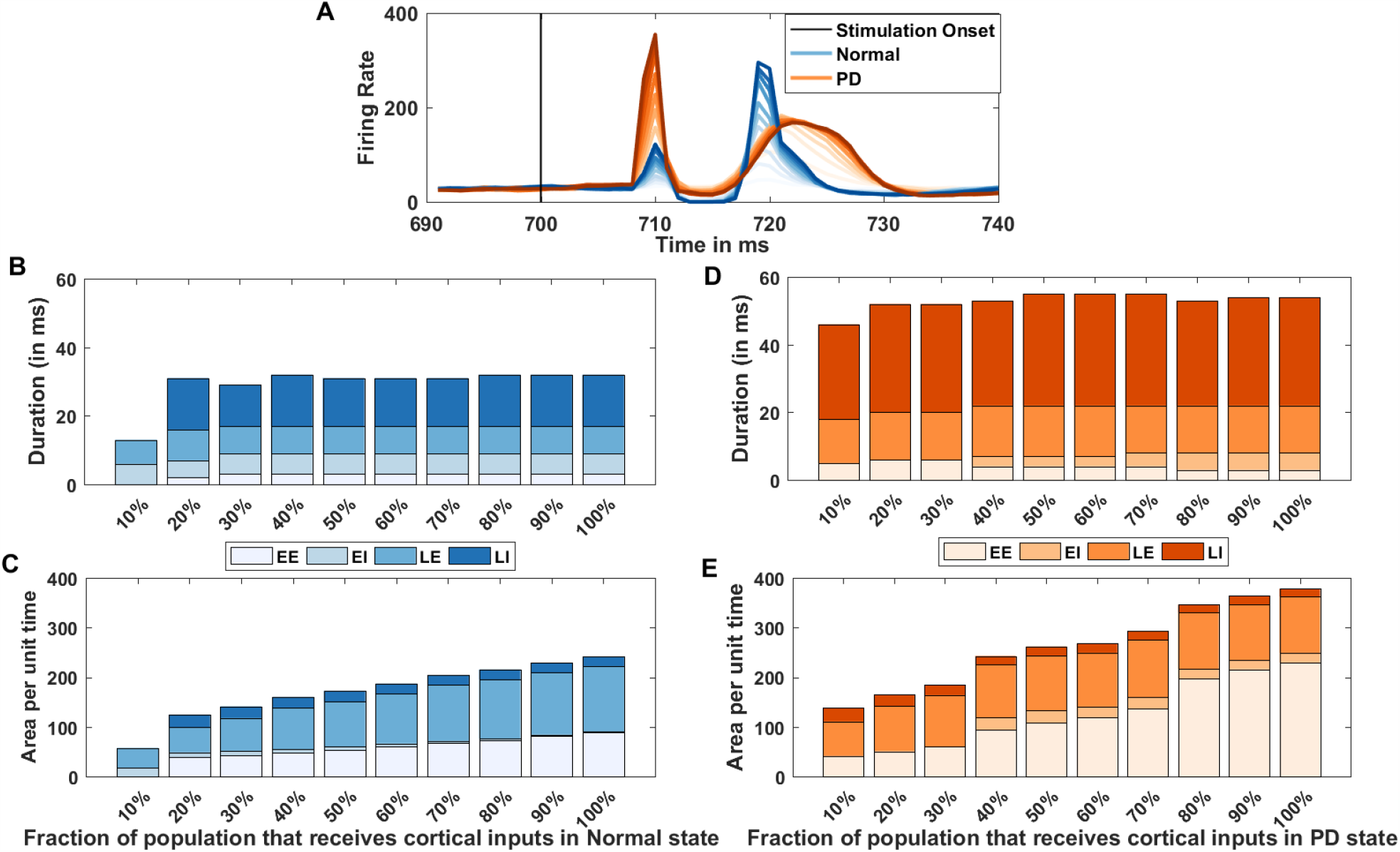
Effect of strength of cortical stimulation on BG transient response shape. To vary the strength of cortical stimulation we varied the fraction of striatal and STN populations that received cortical inputs from 10% to 100%. (**A**) Average transient response (100 trials) in SNr in normal (blue color) and PD state (orange color). Lighter (darker) colors-shades indicate smaller (larger) size of stimulated population. Note that in PD condition, even the strongest cortical input failed to elicit a response similar to that seen in healthy state. (**B**) Changes in the duration of the four zones of the transient response in normal state. (**C**) Changes in the area per unit time (area/time) of the four zones of the transient response in normal state. (**D**) Same as in **B** but for PD state when the network responded with a triphasic response. (**E**) Same as in **C** but for PD state when the network responded with a triphasic response. Note missing colors in a given bar implies that we could detect the corresponding zone.

We found that in both healthy and PD conditions, the amplitude (Figure 4 A) of the four zones are monotonically increasing before saturating to a maximum value. On the other hand, are per unit time of the excitatory zones are increasing indicating the increase in the excitatory strength, whereas decreasing area/time values in inhibitory zones depict stronger inhibition. Interestingly, for weak cortical stimulation (10%) early excitation was below detection threshold in healthy condition (Figure 4 B, C) but in PD condition (Figure 4 D, E), the same weak stimulation elicited a strong early excitatory response.

Overall, these results show that even with the strongest stimulation in PD condition, we could not reproduce the triphasic response properties observed in healthy state with the weakest cortical stimulation. This suggests that the differences in the transient response are not simply due to the altered cortico-BG projections but are primarily because of the altered connectivity within the BG.

### Effect of change in Synaptic connections on cortical evoked transient response in SNr

In the above we demonstrated the existence of a triphasic response pattern for a single combination of synaptic strengths. The total space of different synaptic parameters is of 22-dimension (Table 2) and therefore it is not feasible to test the robustness of our results in a systematic manner by varying all the connection parameters. The structure of BG connectivity suggests that the triphasic response pattern is shaped by D1-SPN→SNr (early inhibition), GPe-TA↔GPe-TI and STN ↔GPe-TI (late excitation/inhibition) connectivity. Therefore, we individually varied these six connections and quantified the duration and area per unit time (area/time) of the four zones of the triphasic response. The minimum and maximum values of each of the synaptic weight (except D1-SPN→SNr synapses) corresponded to their values in l-dopa induced dyskinesia (LID) and PD conditions, respectively. For the case of D1-SPN*→*SNr synapses minimum and maximum values corresponded to PD condition and LID, respectively (see Table 10). If in PD condition, value of a synaptic weight was changed by a factor *m* ∈ ℝ^+^ with respect to its value (*v*) used in the normal condition, then the value of this synaptic weight was varied from v*m to v/m in seven steps.

We found that the duration of the four zones of the triphasic response are robust to changes in these six different synaptic weights (Figure 5 A-F). By contrast, the area/time of the the four zones was sensitive to synaptic weight changes. For instance, the D1-SPN→SNr connection affected the area/time of the EI zone (Figure 5 G); the STN→GPe-TI connection affected the area/time of the EE and EI zones (Figure 5 K) and the D2-SPN→GPe-TI connection affected the area/time of the EI, LE and LI zones (Figure 5 H). However, connectivity between GPe-TI↔GPe-TA and GPe-TI→STN did not affect the area/time of any of the zones (Figure 5 I,J,L).

**Figure 5:**
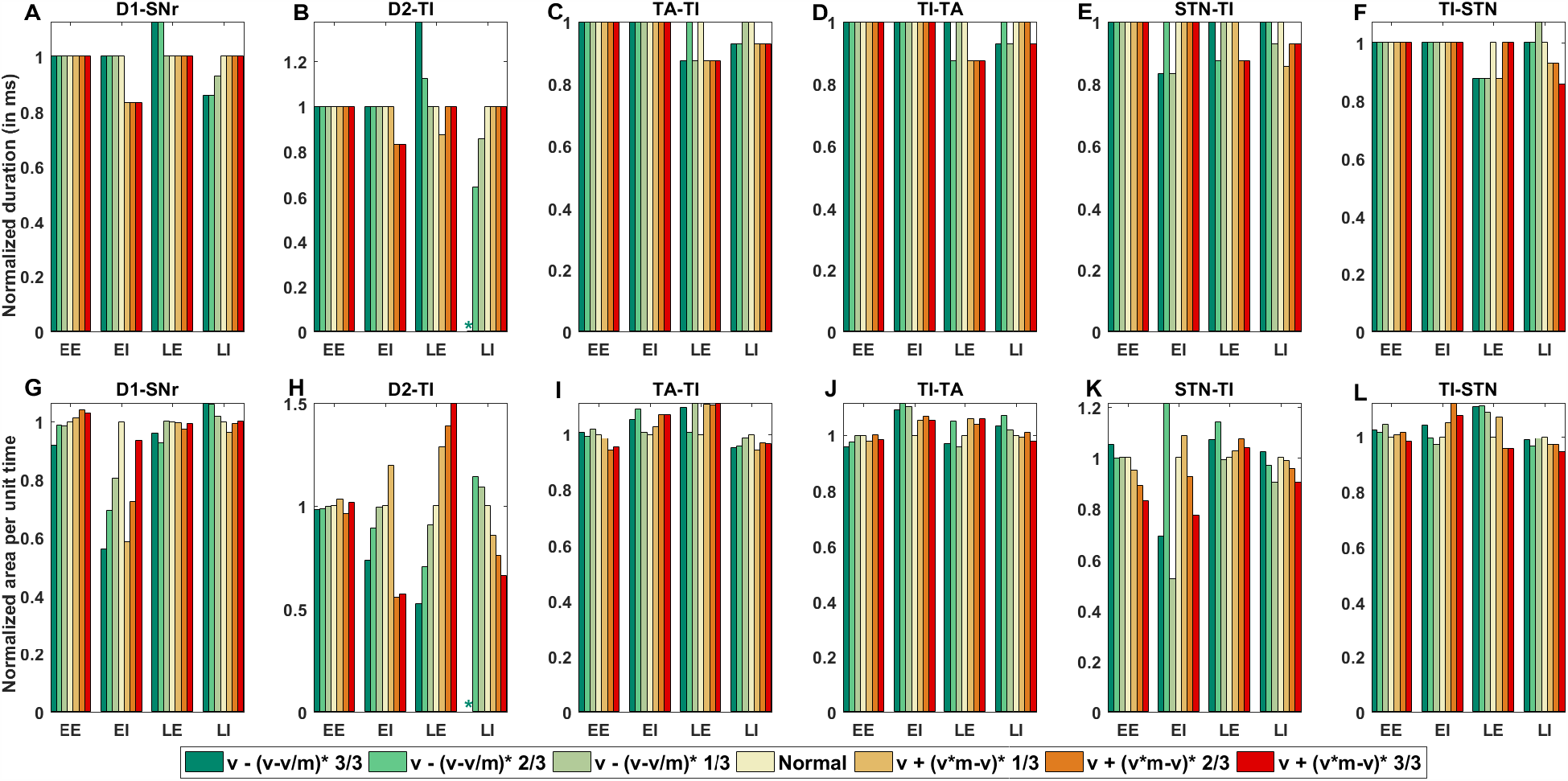
Effect of different synaptic weight changes on SNr activities in normal condition. (**A-F**) Variation in zone wise (normalized) duration for different synaptic connections. Values of zone wise (normalized) duration were normalized with respect to duration in the normal condition for that particular zone. (**G-L**) Variation in zone wise (normalized) area per unit time (area/time) for different synaptic connections. Values of zone wise area per unit time (area/time) were normalized with respect to area per unit time in the normal condition for that particular zone. *v, v*/*m* and *v* * *m* (where *m* ∈ ℝ^+^) denote the value of a particular synaptic strength in normal, high (LID: greenish colors), and low dopamine state (PD: reddish colors), respectively. LI: Late Inhibition, LE: Late Excitation, EI: Early Inhibition, EE: Early Excitation, *: Not Detected

From this analysis, the D2-SPN→GPe-TI emerged as a the most crucial parameter in shaping the transient response in both low and high-dopamine conditions. For extreme values of D2-SPN→GPe-TI connection, area/time of late excitation was very high in low dopamine condition and late inhibition zone was completely absent in high dopamine state.

### Effect of restoration of dopaminergic synaptic connection on the transient response in SNr

In order to simulate PD condition we altered several connectivity parameters (see Table 10). However, in the previous section we showed that the D1-SPN→SNr and D2-SPN→GPe-TI have the strongest effect on the triphasic response. Therefore, next we asked a question red: if we restore selected connections along the indirect pathway (i.e. D2-SPN→GPe-TI or STN↔GPe-TI loop or GPe-TA↔GPe-TI), could we restore the shape of triphasic response to the shape observed in healthy condition?

To this end, first we tuned the BG network in the PD state (Table 10) such that the SNr shows a triphasic response. Then restored the strength of D2-SPN→TI, STN ↔GPe-TI loop, GPe-TA↔GPe-TI, and D1-SPN→SNr one by one. During the restoration of the synaptic connection between a pair of nuclei, (i) the synaptic weights and delays were made equal to normal, (ii) the basal firing rates were made similar to the normal by changing the background firing rate or the background current, (iii) basal firing of SNr was kept the same as that of the normal condition. To compare the triphasic response in healthy and PD states (with and without restoration of certain synaptic weights) we measured the distance between two network conditions using equation 5 (See the subsection **Data Analysis**).

We found that restoring either the D2-SPN→GPe-TI or GPe-TI↔GPe-TA alone can bring the shape of the transient response close to the one observed in healthy state (Figure 6). By contrast, restoration of the D1-SPN→SNr synaptic connection made the network activities different from as in both PD and healthy conditions (Figure 6). This is because in PD condition, in addition to the weakening of D1-SPN to→SNr synapses, cortical inputs to D1-SPN were also weakened (Lindahl and Kotaleski, 2016) and therefore, cortical stimulation only evokes a weak response in D1-SPN. Thus we observed that even though restoration of the D2-SPN→GPe-TI or GPe-TI↔GPe-TA makes the transient response in PD condition more similar to the healthy condition, but even with that the early inhibitory phase is not restored.

**Figure 6:**
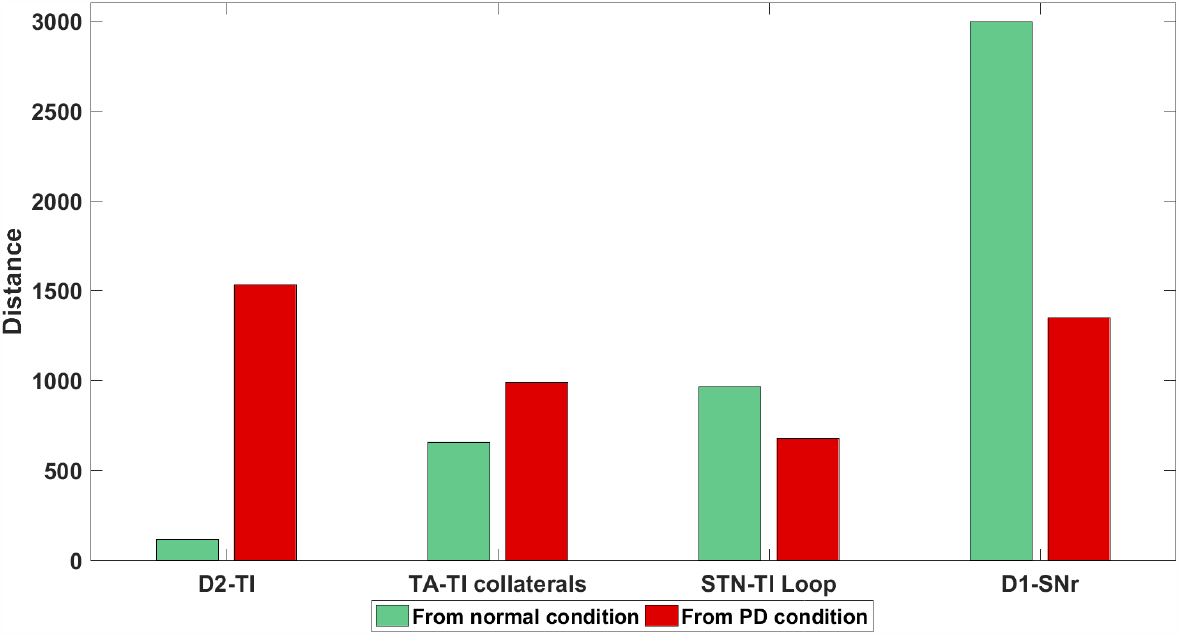
Effect of restoration of synaptic weights between D1-SNr, D2-TI, STN-TI loop, and GPe-TA-GPe-TI with collaterals after dopamine depletion. The green bars indicate the distance (calculated using equation 5) between healthy network and test network whereas red bars indicate the distance between PD network and test network. Here the test network refers to a PD network (tuned to generate triphasic response) in which individual synaptic weights (mentioned on the x-axis) were restored to their healthy value.

### Ongoing activity of BG network

The *β*-band oscillations and synchrony within and between different BG subnuclei in the ongoing activity state (stimulus free) are two prominent hallmarks of PD condition *in vivo* (Brown et al., 2001; Mallet et al., 2008, 2006). Therefore, next we tested whether the network parameters we used to generate the aberrant triphasic and biphasic responses could also induce *β*-band oscillations. To this end, we tuned the BG network in PD condition when it showed either triphasic or biphasic transient response and measured the oscillations and synchrony in the ongoing (stimulus free) activity.

We found that indeed the same set of parameters that generated aberrant transient responses were sufficient to elicit clear *β*-band oscillations in both the biphasic and triphasic response modes (Figure 7 A-D). Next, we measured the phase relationship between different subnuclei of the BG. Mallet et al. (2008) reported that there exists an in-phase relationship between activities of GPe-TA and STN neurons and anti-phase relationship between GPe-TA and GPe-TI neurons. In our model the phase relationships between GPe-TA and Gpe-TI, GPe-TA and STN, GPe-TI and STN (Figure 7 E-G) were similar to that observed in experimental data. Thus, these results suggest that similar changes in the network connection could underlie the aberrant transient response and ongoing activity in PD conditions as well.

**Figure 7:**
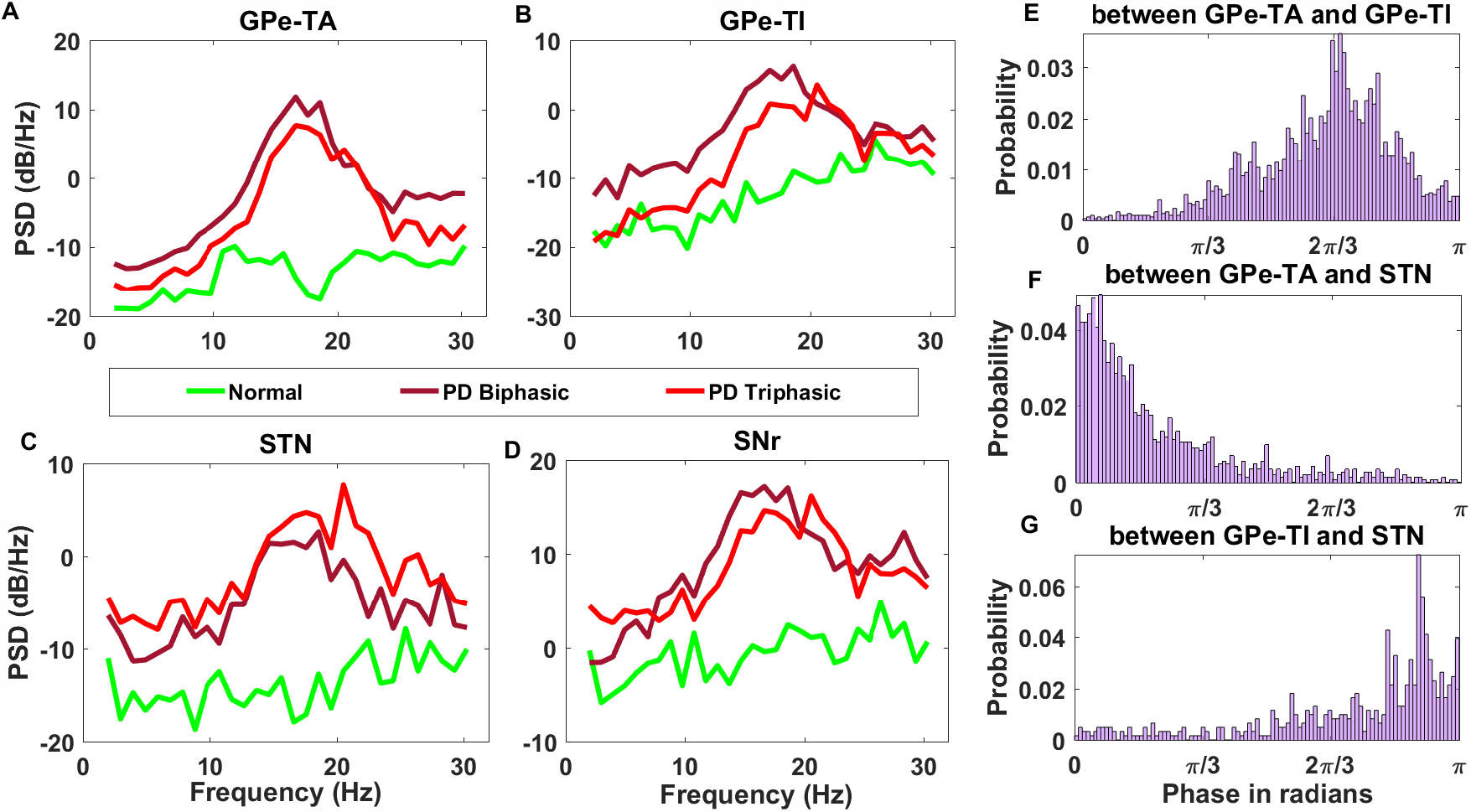
*β*-band oscillations in the ongoing activity of the BG. (**A**) Spectrum of the GPe-TA activity in healthy (green), PD-triphasic (red) and PD-biphasic (brown) response conditions. (**B**) Same as in panel **A** but for GPe-TI. (**C**) Same as in panel **A** but for STN. (**D**) Same as in panel **A** but for SNr. (**E**) Phase relation between GPe-TA and GPe-TI shown in the range of 0 and *π* (in radians). (**F**) Same as **E** but for the phase relation between GPe-TA and STN. (**G**) Same as **E** but for the phase relation between GPe-TI and STN. The phase difference between GPe-TA and GPe-TI attains a peak around 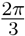 in the phase histogram shown in **E**. The in-phase relation between GPe-TA-STN and approximate anti-phase relation between GPe-TI-STN can be seen in **F** and **G**, respectively.

### Effect of Striato-Pallaidal and Pallido-Subthalamic pathways on the *β*-Oscillations

While *β*-band oscillations are a clear neural signature of PD, the mechanisms underlying the emergence of these oscillations are still not understood. Both experimental data (Plenz and Kital, 1999; Hammond et al., 2007; Tachibana et al., 2011; de la Crompe et al., 2020) and computational models (Kumar et al., 2008; Tachibana et al., 2011; Holgado et al., 2010; Pavlides et al., 2015; Bahuguna et al., 2020) have implicated essentially all the various network interactions in generating oscillations. Here we have developed the BG model primarily to understand the transient response and found that the same model can also generate *β*-band oscillations. Thus, we have a more constrained model of the BG than used previously and this could help us narrow down on the key determinants of oscillations.

Based on our simulations and available experimental data GPe has emerged as a key network necessary to induce *β*-band oscillations. However, it remains unclear which of its input and output connections are more crucial to generate oscillatory activity. Therefore, to quantify the relative contribution of GPe connectivity we either removed striatal input to GPe-TI neurons, or GPe feedback to the striatal FSIs, or GPe-STN interactions. All these perturbations were performed in two different BG networks which showed biphasic or triphasic response in PD condition. In both PD conditions (biphasic response and triphasic response) removal of D2-SPN input to GPe-TI neurons reduced the oscillations and synchrony to nearly to a level observed in healthy state (Figure 8 pale green bars). This supports the hypothesis that increase in D2-SPN activity in dopamine depleted state is responsible for unleashing oscillations in the BG (Kumar et al., 2008; Mallet et al., 2006; Sharott et al., 2017). Moreover, recent experiments also suggest that D2-SPN inputs control oscillations in the GPe-TI population (de la Crompe et al., 2020).

**Figure 8:**
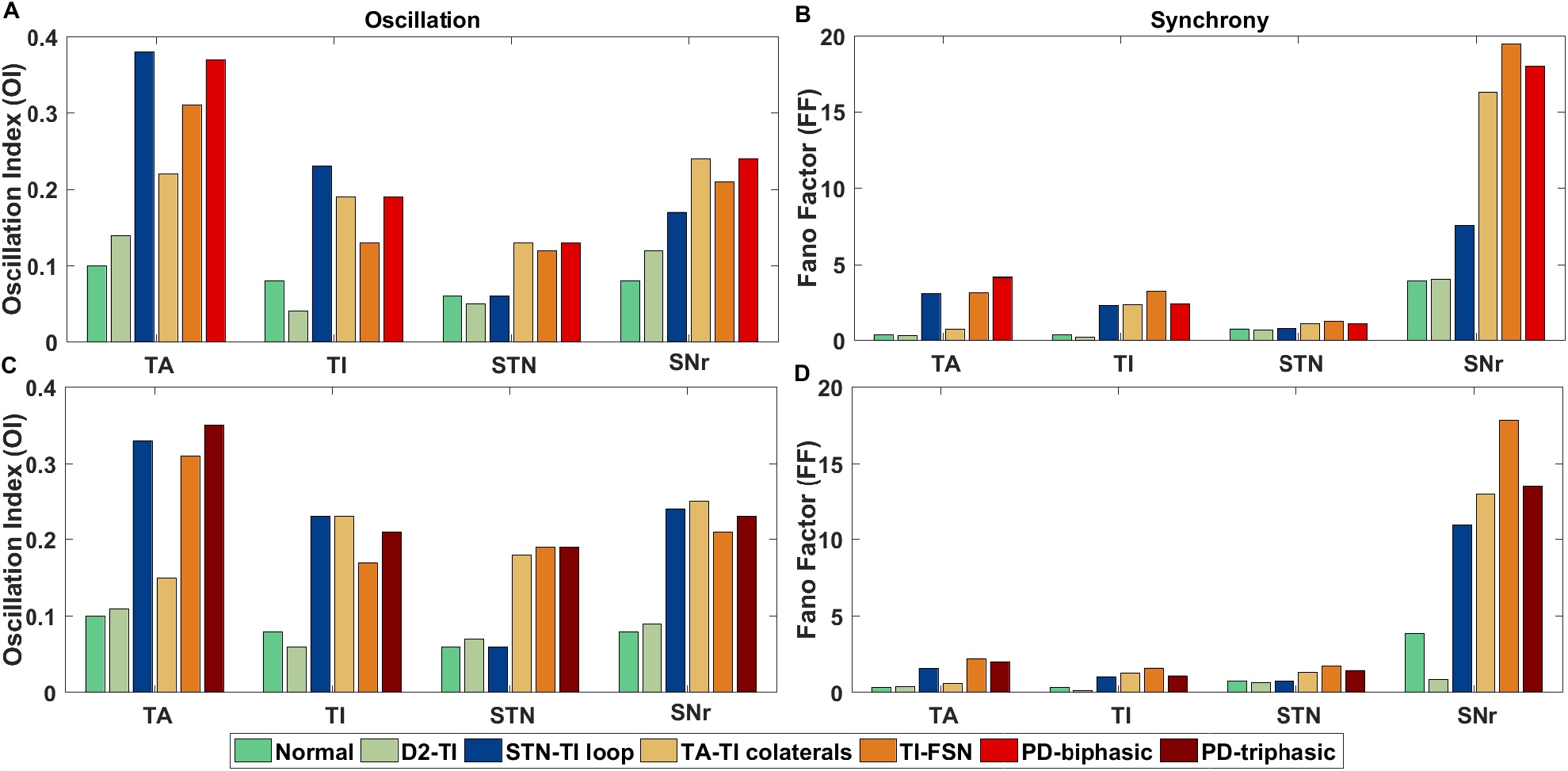
Comparison of relative changes in oscillation and synchrony. (**A**) Oscillation Index for GPe-TA, GPe-TI, STN and SNr with normal, PD-biphasic states and with lesioned networks when synaptic connections between D2-SPN→Gpe-TI, STN↔GPe-TI loop, GPe-TI↔GPe-TA collaterals and GPe-TI →FSI were disconnected. (**B**) Fano Factor for GPe-TA, GPe-TI, STN and SNr with normal, PD-biphasic states and with lesioned networks when synaptic connections between D2-SPN→Gpe-TI, STN↔GPe-TI loop, GPe-TI↔GPe-TA collaterals and GPe-TI→FSI were disconnected. (**C**) Same as **A** while comparing with the PD-triphasic state. (**D**) Same as **B** while comparing with the PD-triphasic state.

By contrast, removal of GPe feedback to striatal FSIs did not affect oscillations or synchrony by much. The effect of interactions within and between GPe-TA and GPe-TI neurons was dependent on the state of the network: removal of these connections reduced oscillations and synchrony by a larger amount when the BG was tuned to exhibit triphasic response in PD condition (Figure 8 pale orange bars). However, in even after removal of collateral within the GPe neurons, both oscillations and synchrony were much higher than that observed in a healthy state.

Surprisingly, removal of STN↔GPe-TI connections did not affect the oscillation, irrespective of the type of transient responses the network showed in PD condition (Figure 8 blue bars). These findings imply that removal or inhibition of the STN will also not have any effect on the oscillations. This is consistent with the recent findings by de la Crompe et al. (2020) who showed that optogenetic inhibition of STN does not quench oscillations (see also Gradinaru et al. (2009)).

### Diversity of transient responses

As noted earlier, our BG network model is homogeneous and therefore, we could either generate biphasic or triphasic shaped transient response in the network. This approach however, allowed us to identify the key network interactions that are involved in changing the response shape from biphasic to triphasic (i.e. D2-SPN →GPe-TI, D1-SPN→SNr, and GPe-TI→STN). An inhomogeneous change in these connections could be one reason for the observed diversity of transient responses in *in vivo*. However, oscillations in the ongoing activity could also contribute to the diversity of transient responses because the shape of transient response may depend on the oscillation phase at which cortical stimulation was delivered. In fact, recent experimental data suggests that when *β* band oscillations are weak or absent in PD, transient responses variability is reduced (Chiken et al., 2020).

To test this hypothesis, we tuned the network in a PD state in which it responded with a biphasic shape (Table 10) and delivered the stimulus at different phases of oscillations. We pooled the data and observed six types of responses namely, “EE-EI-LE”, “EE-EI”, “EI-LE”, “EE-LE”, “EE” and “LE”. Here “EE-EI-LE” denotes a triphasic response consisting of early excitation (“EE”), inhibition (“EI”) and late excitation (“LE”). The other responses were variations of that, where a partial response was seen in each of them. The relative fraction of each of the six responses is shown in Figure 9 A. Because the network was tuned to generate biphasic responses, the majority of the responses (47%) happened to be “EE-LE” (similar to Figure 3 B). However, we also observed monophasic and triphasic responses as well (Figure 9 A).

**Figure 9:**
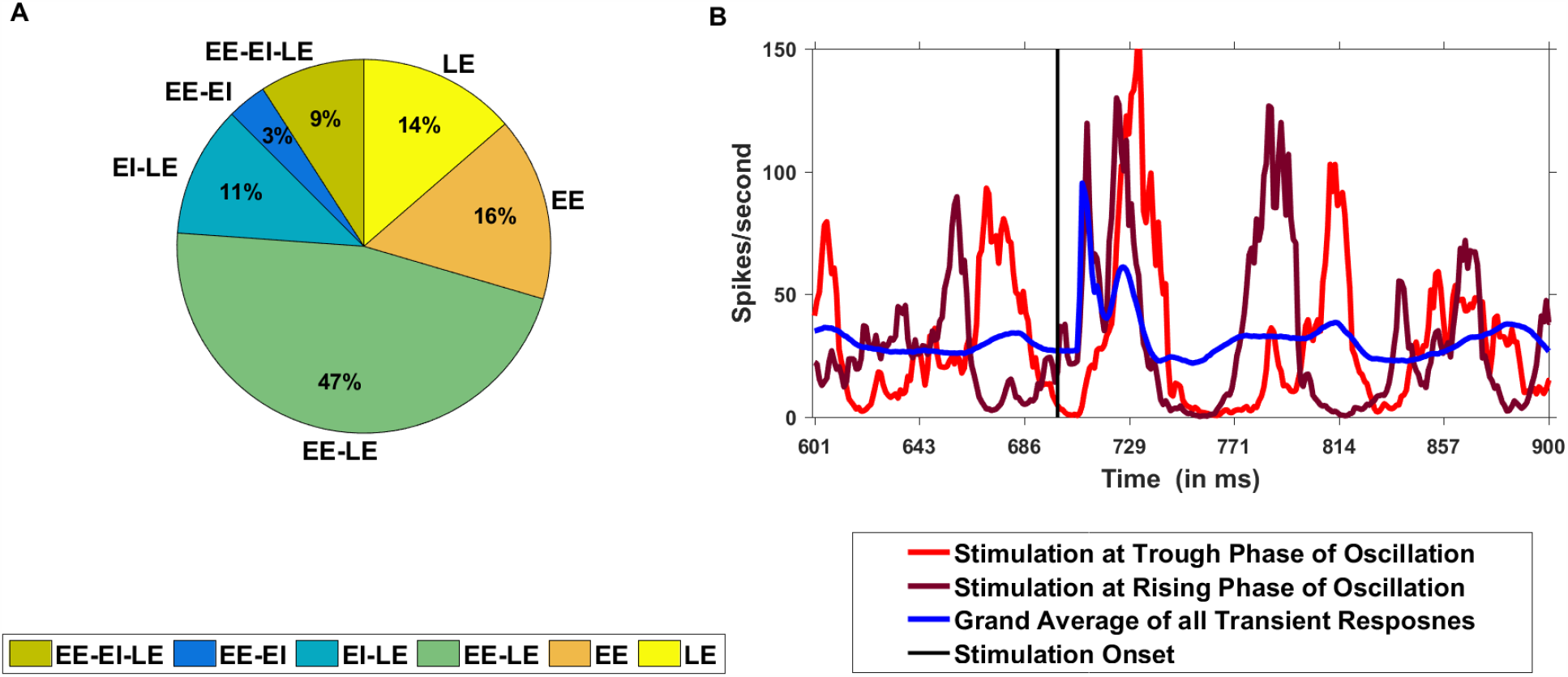
Diversity of transient response may depend on the phase of cortical stimulation. (**A**) Percentage of six types of SNr responses namely, EE-EI-LE, EE-EI, EI-LE, EE-LE, EE and LE. (**B**) The shape of transient response depends on the oscillation phase. Blue trace: average transient response (4224 trials) obtained by stimulating the BG at random phases (88 in number to cover the full 2*π* phase in the SNr) of oscillation. Red trace: Average transient response (48 trials), categorized as “LE” when the stimulation arrived at the trough of oscillation in SNr. Brown trace: average transient response (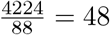 trials), categorized as “EE-LE” when the cortical input arrived at a rising phase of the oscillation in SNr.

This variation was primarily due to the differences in phase of the oscillation at which cortical stimulation was delivered. For instance, when the stimulus arrived at the falling edge close to the trough of the *β*-oscillations (corresponding to a phase delay of 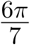 from the previous peak), SNr was not able to respond to the hyper-direct pathway and the EE was not visible (red trace, Figure 9 B). Therefore, these types of responses were observed as “LE”. On the contrary, when the stimulation arrived at the rising phase of the oscillation (corresponding to a phase delay of 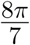 from the previous peak), both early and late excitation were visible, whereas the inhibition was not strong enough (brown trace, Figure 9 B).

In order to characterize the contribution of different BG nuclei to the transient response in healthy and PD conditions, we varied the strength of several connections in BG (e.g. see Figure 4,5). We pooled all those simulations together, where the synaptic connection strengths were varied according to Figure 5, and estimated the variability of the transient responses. The rationale to do this was that each network simulation with different connection strength may represent a different SNr/GPi region or animal where the transient response was recorded. Indeed, such pooling of the data resulted in a high heterogeneity in the transient responses in both healthy and PD conditions (see Table 12) which closely matched with the experimental data. These results while do not explain the full diversity of the responses observed in *in vivo*, they show that the oscillation phase as well as diversity of synaptic connectivity are important variables in determining the shape of the response.

**Table 12:** Features of the transient response of the SNr neurons, same as Table 11 however, by pooling synaptic weights corresponding to Figure 5. Here the variations in the features in normal state were obtained by simulating the network with the range of synaptic weights of a particular connection between v - (v - v/m)*1/3 and v + (v*m - v)*1/3. Similarly, the variations in the features in the PD conditions (PD-biphasic and PD-triphasic) were obtained by simulating the network with the range of synaptic weights of a particular connection between v + (v*m - v)*1/3 and v + (v*m - v)*3/3. These were done by considering 6 types of synaptic connections corresponding to Figure 5.

|  | Normal (triphasic) | PD-biphasic | PD-triphasic |
| --- | --- | --- | --- |
| **Early Excitation (EE)** |  |  |  |
| Latency (ms) | $7.0 \pm 0$ | $7.0 \pm 0$ | $7.0 \pm 0$ |
| Duration (ms) | $3.0 \pm 0$ | $23.0 \pm 0.79$ | $3.69 \pm 0.46$ |
| Deviation of peak amplitude from the baseline | $38.02 \pm 2.76$ | $75.84 \pm 12.53$ | $153.63 \pm 6.43$ |
| **Early inhibition (EI)** |  | *ND |  |
| Latency (ms) | $10.0 \pm 0$ | | $10.69 \pm 0.46$ |
| Duration (ms) | $5.56 \pm 0.49$ | | $3.66 \pm 0.77$ |
| Deviation of peak amplitude from the baseline ($H_p - H_{bas}$) | $-29.18 \pm 0.54$ | | $-11.7 \pm 1.73$ |
| **late excitation (LE)** |  | ## |  |
| Latency (ms) | $15.56 \pm 0.49$ | | $14.35 \pm 0.47$ |
| Duration (ms) | $7.66 \pm 0.51$ | | $14.21 \pm 0.71$ |
| Deviation of peak amplitude from the baseline ($H_p - H_{bas}$) | $156.72 \pm 16.24$ | | $139.45 \pm 2.91$ |
| **Late inhibition (LI)** |  |  |  |
| Latency (ms) | $23.22 \pm 0.42$ | $30.0 \pm 0.79$ | $28.57 \pm 0.49$ |
| Duration (ms) | $13.17 \pm 0.96$ | $44.09 \pm 9.35$ | $31.8 \pm 1.11$ |
| Deviation of peak amplitude from the baseline ($H_p - H_{bas}$) | $-11.81 \pm 1.64$ | $-16.6 \text{ inhibition} \pm 1.72$ | $-19.55 \pm 1.31$ |

## Discussion

Here we have studied how the changes induced by low-dopamine affect both transient response (induced by cortical stimulation) as well as the ongoing activity state of the BG network. Typically, a transient stimulation of the cortex results in a triphasic response in the SNr/GPi (the output of the BG). The shape of the response is impaired in chronic low-dopamine conditions such as Parkinson’s disease. The different zones of the transient response can be associated with different aspects of initiation of voluntary movements. For instance, it has been hypothesized that EE zone resets the cortical activity, EI zone allows for the execution of movements and LE zone stops the movement (Nambu et al., 2002; Chiken et al., 2020). A weaker or completely absent EI zone in PD is thought to be related to akinesia. Indeed, L-dopa treatment or local inhibition of the STN both of which restore the EI zone also ameliorate motor deficits in PD (Chiken et al., 2020). The triphasic response in the SNr/GPi is usually explained by difference in the relative timing of the direct, indirect and hyper-direct pathways of the BG which converge in the SNr/GPi.

Here, we show that changes in the shape of the transient response in PD state involve not only changes in the feed-forward connections between different subnuclei of the BG (D1-SPN→SNr) but also by interactions between STN and GPe (GPe-TI↔STN) (Figure 5 K) and to some extent by GPe-TA↔GPe-TI (Figure 6). Moreover, we show that same changes in the BG network (both synaptic and neuronal excitability) may underlie the impairment of transient response and emergence of induced population level oscillations and synchrony in the BG.

In PD condition, neurons either show biphasic or triphasic transient responses (Sano and Nambu, 2019), the later is however quantitatively different from the triphasic response observed in healthy state. In our model, the aberrant biphasic response in PD condition appeared as we changed the parameters to a low-dopamine state (according to the model by Lindahl and Kotaleski (2016)). However, to obtain a triphasic response, we needed to reduce D2-SPN→GPe-TI, increase D1-SPN→SNr and reduce GPe-TI→STN connections (see Table 10). This suggests that dopamine effects are not homogeneous within and between different subnuclei of BG. To restore a healthy state it is important to experimentally characterize the heterogeneity of dopamine action. The diversity of dopamine action and phase of oscillations at which stimulation was delivered, together could explain the observed diversity of transient responses in *in vivo*.

Previously, Blenkinsop et al. (2017) suggested that in a healthy state, biphasic and triphasic responses in the SNr arise because of interactions among functionally segregated channels of competing inputs with different strengths. In a BG model with functionally segregated channels, local inhibition within the GPe and excitation from a small number of highly active STN neurons (presumably because of stronger cortical inputs) are responsible for emergence of LE zone, rendering a response biphasic or triphasic (Blenkinsop et al., 2017). Here, we have used a BG model without functionally segregated channels. Our results suggest that diversity of synaptic strengths within and between BG nuclei could give rise to some neurons responding in a triphasic and others in a biphasic manner. Consistent with the model by Blenkinsop et al. (2017) in our model the magnitude of LE zone can be controlled by the strength of cortical stimulation (Figure 4). Our work points out a strong influence of indirect pathway (D2-SPN to GPe-TI) in controlling the shape of the transient response both in normal and PD condition. This observation is also consistent with the proposal of Blenkinsop et al. (2017).

Even though we managed to generate a triphasic response in PD condition, it was quantitatively different from that one observed in a healthy state. The differences were most clearly seen in the late excitation which was weaker in amplitude, but lasted longer in PD condition as compared to a healthy state. Moreover, these differences in the triphasic response could not be compensated by increasing the magnitude of the cortical stimulation, suggesting that impaired transient response also entails impaired recurrent interactions within and between BG subnuclei.

Here we assumed that the GPe to SNr and STN to SNr synapses are static. However, experimental data suggests that synapses between GPe to SNr show short-term depression (Connelly et al., 2010). Lindahl et al. (2013) has argued that when GPe to SNr synapses show short-term depression, the STN to SNr synapse should also show short-term depression in order to keep the SNr response small. Lindahl et al. (2013) further showed that short-term depression can have a big effect on the response of the BG when inputs last 100s of milliseconds. Here, in our model we have only considered very short-lasting stimuli and therefore, short-term depression of synapses might not affect our results. However, this should be tested in a more detailed model.

Dopamine has multiple effects on neuron’s excitability, synaptic strength, and synaptic plasticity (see table 10). To better understand which one of these are most detrimental for the shape of the transient response, we individually perturbed six of the most crucial parameters (Figure 5). This analysis revealed that the connection D2-SPN→GPe-TI is the most crucial for the shape of the transient response as it controls both late excitation and late inhibition zones (Figure 5). In addition, D1-SPN→SNr connection is expected to be crucial for determining the early inhibition zone. We further corroborated these results by restoring the strength of D2-SPN→GPe-TI connection to their normal level while keeping all other parameters to their low-dopamine levels. This single change was effective in bringing the triphasic response in PD state closer to the one observed in healthy state.

In network models of *β*-band oscillations when both STN and GPe are included, invariably STN↔GPe connections emerge as a key parameter in shaping the oscillations (Holgado et al., 2010; Pavlides et al., 2015). In the full model of BG with both striatum and cortico-BG loop, STN↔GPe may not be as important. Indeed Leblois et al. (2006) showed that altered interactions among direct and hyper-direct pathways are sufficient to induce oscillations. However, in the model by Leblois et al. (2006) GPe plays no role in generating oscillations – this is inconsistent with the experimental data (de la Crompe et al., 2020). In our model, consistent with the recent experimental data (de la Crompe et al., 2020) STN↔ GPe is not important for generating oscillations. In fact, in our model removal of STN↔GPe-TI connections did not affect the oscillations (Figure 8 blue bars). These observations combined with the experimental data (de la Crompe et al., 2020) raise the question which network interactions generate oscillations, if not the STN-GPe loop. We have not explored this question in this work as the question requires a more systematic study. However, we speculate that besides the STN-GPe, the back-projections from GPe to striatum together with recurrent connections within the GPe can form an effective excitatory-inhibitory network necessary for generating oscillations. It is worth noting that previous experimental data (Mallet et al., 2006; de la Crompe et al., 2020; Sharott et al., 2017) and computational models (Kumar et al., 2011; Mirzaei et al., 2017; Bahuguna et al., 2020) provide a strong evidence that strengthening of D2-SPN→GPe-TI connection is also sufficient to induce *β*-band oscillations/synchrony in the ongoing activity state of the BG. Thus, here, we provide a unified explanation of impaired transient response and ongoing activity in PD state. Our results highlight the importance of the GPe in controlling the dynamics and function of the BG.

Despite its simplicity, our model not only provides network interaction that shapes the properties of transient responses in the BG, but also the model clearly suggests that recurrent interactions within and between subnuclei of BG are crucial in shaping the transient response. We found that the duration of different zones of the transient responses is largely robust to changes in the BG network interactions while the area/time of the different zones is not. This implies that in *in vivo* data we should find a narrow distribution of the duration of different zones and a wider distribution of the area/time of different zones. Next, our model predicts that by strengthening of cortical inputs, the normal shape of transient response cannot be restored in PD state. This prediction can be tested by either increasing the stimulus strength or by increasing the number of stimulated neurons (e.g. using optogenetic stimulation methods). Finally, the model predicts that by restoring the normal strength of D2-SPN→GPe-TI (or also by reducing the activity of D2-SPN), a near healthy shape of transient response could be restored even in PD condition.

## Acknowledgements

We thank Dr. Jyotika Bahuguna for helpful discussions and feedback on the manuscript. The author AK acknowledges funding from Swedish Research Council (Grant VR-M-2018-03118), StratNEURO, KTH Digital Futures project dBRAIN and STINT (Joint Japan-Sweden Research Collaboration). The author JHK acknowldged funding from Swedish Research Council (Grant VR-M-2017-02806) KTH digital futures project dBRAIN, Swedish e-Science Research Center, European Union Seventh Framework Programme (Grant FP7/2007-2013), EU/Horizon 2020 (Grants 720270 HBP SGA1; Grant 785907 HBP SGA2), European Union/Horizon 2020 (Grant no. 945539, Human Brain Project, SGA3). The authors KC, AS, SR acknowledge funding from TCS Research & Innovation division of the Tata Consultancy Services (to KC, AS, SR). AN acknowledges MEXT KAKENHI (“Non-linear Neurooscillology”, 15H05873), JSPS KAKENHI (19KK0193) and STINT (Joint Japan-Sweden Research Collaboration). Parts of the simulations were performed on resources provided by the Swedish National Infrastructure for Computing (SNIC) at the PDC Center for High Performance Computing, KTH Royal Institute of Technology.

